# Eicosanoids in the pancreatic tumor microenvironment – a multicellular, multifaceted progression

**DOI:** 10.1101/2021.10.27.466097

**Authors:** Vikas B. Gubbala, Nidhi Jytosana, Vincent Q. Trinh, H. Carlo Maurer, Razia F. Naeem, Nikki K. Lytle, Zhibo Ma, Steven Zhao, Wei Lin, Haiyong Han, Yu Shi, Tony Hunter, Pankaj K. Singh, Kenneth P. Olive, Marcus C.B. Tan, Susan M. Kaech, Geoffrey M. Wahl, Kathleen E. DelGiorno

## Abstract

Eicosanoids, oxidized fatty acids that serve as cell-signaling molecules, have been broadly implicated in tumorigenesis. To identify eicosanoids relevant to pancreatic tumorigenesis, we profiled normal pancreas and pancreatic ductal adenocarcinoma (PDAC) in mouse models and patient samples using mass spectrometry. We interrogated RNA sequencing datasets for eicosanoid synthase or receptor expression. Findings were confirmed by immunostaining. In murine models, we identified elevated levels of PGD_2_, prostacyclin, and thromboxanes in neoplasia while PGE_2_, 12-HHTre, HETEs, and HDoHEs are elevated specifically in tumors. Analysis of scRNA-seq datasets suggests that PGE_2_ and prostacyclins are derived from fibroblasts, PGD_2_ and thromboxanes from myeloid cells, and PGD_2_ and 5-HETE from tuft cells. In patient samples, we identified a transition from PGD_2_ to PGE_2_-producing enzymes in the epithelium during the transition to PDAC, fibroblast/tumor expression of PTGIS, and myeloid/tumor cell expression of TBXAS1. Altogether, our analyses identify key changes in eicosanoid species during pancreatic tumorigenesis and the cell types responsible for their synthesis.

## Introduction

Pancreatic ductal adenocarcinoma (PDAC) can arise from the progression of acinar-to-ductal metaplasia through several grades of pancreatic intraepithelial neoplasia (PanINs), which are characterized by increasing nuclear atypia and loss of cellular polarity. Metaplasia is a process in which acinar cells transdifferentiate into ductal-like cells in response to pancreatic damage to re-establish homeostasis. However, in the context of oncogenic mutations, such as *Kras*^*G12D*^, ADM becomes irreversible and leads to the formation of hyperplastic ductal structures known as PanINs. Accumulation of additional genetic mutations leads to tumor formation (Storz, 2017). Interestingly, ADM does not result in the formation of a homogeneous population of cells. Instead, ADM generates diverse, differentiated, secretory cells, including tuft cells. Tuft cells, or solitary chemosensory cells, have been detected in chronic pancreatitis and *Kras*^*G12D*^-induced tumorigenesis (Delgiorno et al., 2014; DelGiorno et al., 2020b). Recently, we showed that tuft cells attenuate pancreatic tumorigenesis through the synthesis and secretion of the eicosanoid prostaglandin D_2_ (PGD_2_)(DelGiorno et al., 2020a). These cells, which are proposed to play diverse roles in early tumorigenesis and immune cell recruitment, are abundant in preinvasive lesions but are lost in the adenocarcinoma stage of pancreatic tumorigenesis (Delgiorno et al., 2014).

PDAC tissue largely consists of desmoplastic stroma, which supports a diverse population of cell types, including tumor epithelium, immune cells, and other stromal cell types. One such stromal cell type is the cancer-associated fibroblast (CAF), which functions in a variety of roles including ECM deposition, immune signaling, and tumor cell support (Elyada et al., 2019). CAFs themselves are a heterogeneous class of cells and can be further subdivided into cell states with specific roles. These subtypes include inflammatory CAFs (iCAFs), which secrete cytokines and chemokines, myofibroblastic CAFs (myCAFs), which primarily deposit extracellular matrix, and antigen presenting CAFs (apCAFs), which have only been detected as a distinct subset in mouse models of PDAC (Biffi et al., 2019; Elyada et al., 2019). PDAC are also enriched with several immune cell types, including neutrophils, macrophages, dendritic cells, and B- and T-lymphocytes.

Cellular crosstalk within the tumor microenvironment can significantly affect tumor progression. One important class of signaling molecules involved in this crosstalk are eicosanoids, or lipid products derived from the enzymatic or non-enzymatic oxidation of polyunsaturated fatty acids (PUFAs)(Wang and Dubois, 2010). Hundreds of eicosanoids have been characterized, each with distinct, context dependent roles based on the presence and characteristics of receptors expressed in the tumor microenvironment. Eicosanoids can be further subdivided into broad classes – including prostanoids, HETEs, and HdoHEs – based on the precursor molecule PUFA and the synthases responsible for their production.

Arachidonic acid is a common precursor for many different eicosanoids and certain phospholipases (PLA2G2A, PLA2G4A) can generate arachidonic acid from membrane phospholipids. Synthesis of prostanoid-type eicosanoids begins with metabolism of arachidonic acid by cyclooxygenases (COX1, COX2), to prostaglandin G_2_ (PGG_2_), which is non-enzymatically metabolized to prostaglandin H_2_ (PGH_2_). Terminal synthases are required to convert this substrate to specific prostanoid molecules (Figure 1A)(Bui et al., 2011; Wang and Dubois, 2010).

**Figure 1.**
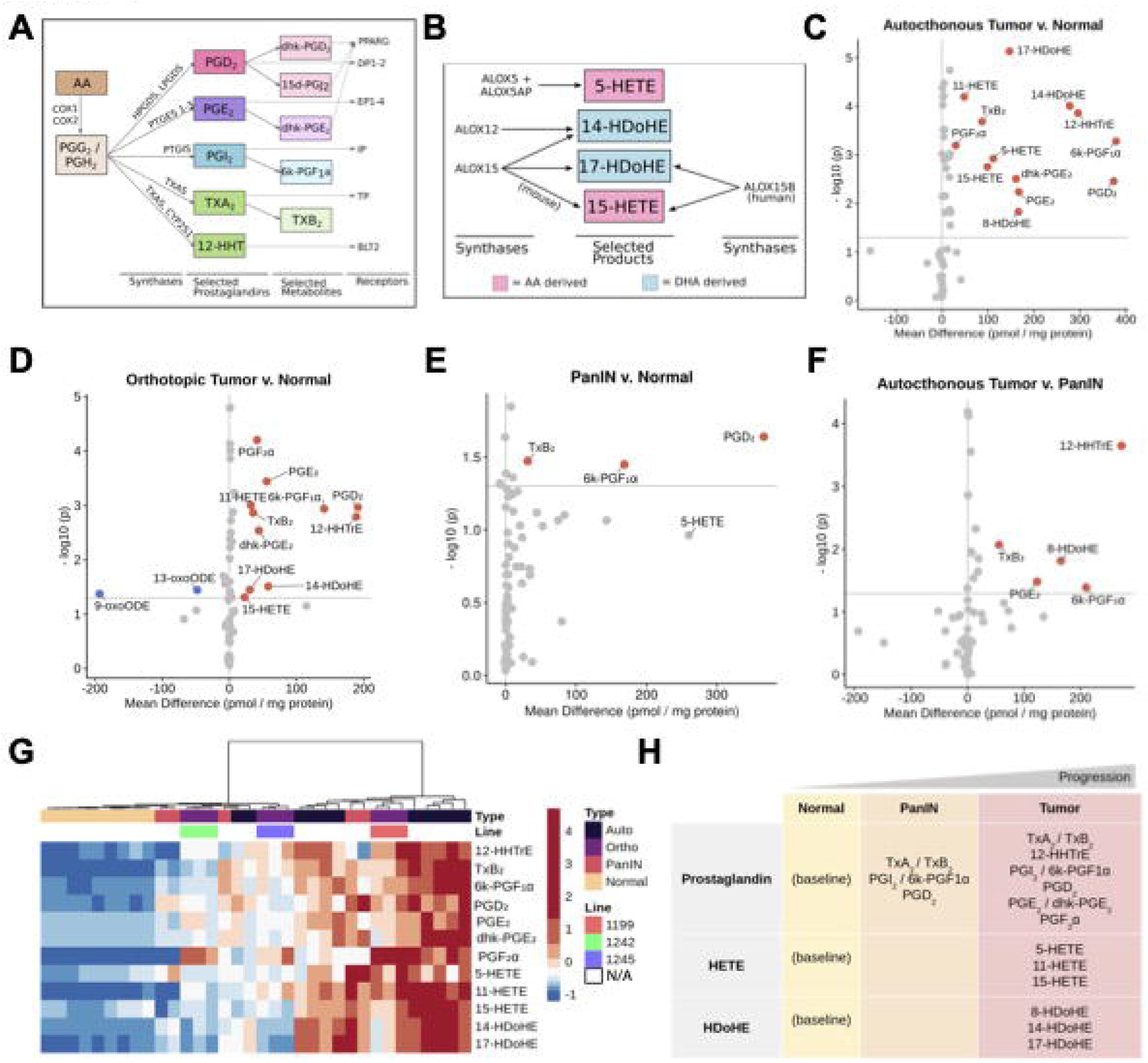
Eicosanoid levels throughout disease progression in mouse models of pancreatic tumorigenesis. Schematics depicting biosynthesis of select **(A)** prostaglandins and **(B)** HETEs/HDHEs, along with relevant synthases and receptors. Eicosanoid profiles of **(C)** autochthonous tumors (*KPC* mice), **(D)** orthotopic tumors (FC-1199, FC-1242, FC-1245 cell lines), and **(E)** PanIN-bearing pancreata (*KC* mice) with respect to normal pancreas values. **(F)** Eicosanoid levels in autochthonous tumors with respect to PanIN. All values, pmol/mg protein. Samples with a mean difference +/− 20 pmol/mg protein and p < 0.05 are indicated in either red, up, or blue, down. **(G)** Hierarchical clustering and heatmap of select eicosanoid levels in the samples shown in C-F. Heatmap values are z-score normalized by row and colors are assigned by quintile scaling. **(H)** Schematic of major eicosanoids associated with pancreatic tumorigenesis in mouse models.

Prostanoids have been studied in the context of tumorigenesis in a variety of organ systems. Prostaglandin E_2_ (PGE_2_) has been linked to tumor growth, metastasis, and fibroblast function in several cancers including PDAC (Markosyan et al., 2019; Wang et al., 2021; Wang and Dubois, 2010). Its metabolite, dhk-PGE_2_, is generated by the enzyme HPGD (also known as 15-PGDH); HPGD ablation has been shown to accelerate tumorigenesis in the intestines and pancreas (Arima et al., 2019; Yan et al., 2009). Prostaglandin D_2_ (PGD_2_) can suppress tumor cell proliferation and metastasis in intestinal adenomas and gastric cancer (Tippin et al., 2014; Zhang et al., 2018). Thromboxane A_2_ (TxA_2_) typically functions as a platelet activator and vasoconstrictor but has also been implicated in promoting tumor cell growth, metastasis, and regulation of neovascularization in breast cancer, lung cancer and more (Ekambaram et al., 2011; Li et al., 2017; Lucotti et al., 2019). TxA_2_ has a very short half-life and quickly metabolizes to thromboxane B_2_ (TxB_2_), a product predicted to have minimal biological activity (Roberts et al., 1977). Produced in equimolar ratios in thromboxane biosynthesis is prostaglandin 12-HHTre, a relatively understudied eicosanoid with only one known receptor, BLT2 (LTB4R2)(Liu et al., 2014). 12-HHTre can also be produced by cytochrome proteins, such as CYP2S1 (Okuno and Yokomizo, 2018). Although prostacyclin (PGI_2_) antagonizes the actions of TxA_2_ and acts as a vasodilator and inhibitor of platelet activation, it too has been implicated in pro-tumorigenic, pro-angiogenic roles in the tumor microenvironment and is associated with poor prognosis in lung, ovarian, and gastric cancers (Dai et al., 2020; Osawa et al., 2012). It also has a short half-life, and readily breaks down into the inert, stable product 6k-PGF1α (Moncada et al., 1976).

Lipoxygenases oxidize arachidonic acid to generate HETEs (Figure 1B)(Conteh et al., 2019; Mozurkewich et al., 2016; Snodgrass and Brune, 2019). 5-HETE is generated from 5-lipoxygenase (ALOX5); ALOX5 inhibition has been shown to slow the growth of cancer cells *in vitro* (Austin Pickens et al., 2019; Avis et al., 2001; Ghosh and Myers, 1998). The role of eicosanoid 15-HETE, on the other hand, is less clear, as it has been implicated in both pro- and anti-tumorigenic roles (O’Flaherty et al., 2013; Shappell et al., 2001). Lipoxygenases can metabolize the PUFA docosahexaenoic acid (DHA) into HDoHEs (14-HDoHE, 17-HDoHE), biosynthetic precursors to maresins and D-resolvins, respectively (Valdes et al., 2017; Zhang et al., 2017). While the action(s) of HDoHEs in cancer are unknown, their metabolic byproducts can attenuate inflammation and wound healing and maresins can inhibit tumor growth (Sulciner et al., 2018).

Broad inhibition of prostanoid synthesis by targeting the upstream synthase COX2 has been shown to reduce tumor growth and metastasis, reverse collagen deposition, sensitize tumors to immunotherapy, normalize the vasculature, and enhance the response to chemotherapy in mouse models (Kirane et al., 2012; Markosyan et al., 2019; Zhang et al., 2019). However, COX2 inhibition blocks the synthesis of several downstream eicosanoids, causing significant off-target effects. To develop better, more specific therapeutics, it is important to characterize eicosanoid diversity in PDAC. To accomplish this, we conducted eicosanoid profiling on normal pancreata and PDAC in mouse models of pancreatic tumorigenesis and human patient samples. We then interrogated published scRNA-seq datasets to identify the cellular source(s) of eicosanoid synthases and receptors and validated key findings at the protein level.

## Results

### Eicosanoid composition evolves throughout pancreatic tumorigenesis

To broadly profile eicosanoid diversity, we used mass spectrometry and assessed a panel of over 157 eicosanoid species in normal pancreata, autochthonous tumors from either *Kras*^*G12D*^;*Trp53*^*R172H*^;*Ptf1a*^*Cre/+*^ or *Kras*^*G12D*^;*Trp53*^*fl/fl*^,*Pdx1-Cre* mice (collectively referred to as KPC) and syngeneic orthotopic tumors generated from 3 separate PDAC cell lines (derived from *KPC* mice) (Figure S1A). We identified significant increases in mean concentration for several eicosanoids in autochthonous *KPC* tumors (n = 11) as compared to normal pancreata (n = 9) (Figure 1C, mean difference > 20 pmol/mg protein, p < 0.05). Most of the elevated species belong to the prostaglandin class of eicosanoids, including PGE_2_ and its metabolite dhk-PGE_2_, PGD_2_, thromboxane synthesis byproducts 12-HHTrE and TxB_2_, and prostacyclin (PGI_2_) metabolite 6k-PGF1α (Figure 1A, 1C). Other eicosanoids identified in *KPC* tumors are lipoxygenase-derived HETEs (5-HETE, 11-HETE, 15-HETE) and docosahexaenoic acid derived HDoHEs (14-HDoHE, 17-HDoHE, 8-HDoHE) (Figure 1B-C). To determine if eicosanoid levels in orthotopically generated PDAC tumors reflect those identified in autochthonous models, we analyzed tumors generated from three different murine PDAC cell lines (FC-1199, FC-1242, and FC-1245, n = 3/line). Interestingly, most eicosanoids identified as elevated in autochthonous models, relative to normal pancreas, were also elevated in orthotopic tumors except for 5-HETE (Figure 1D). Additionally, analysis of orthotopic tumor profiles identified a significant decrease in linoleic acid oxidation products 9- and 13-oxoODE, as compared to normal pancreas, which did not reach significance in autochthonous PDAC tumor profiles (Figure 1D).

Previously, we profiled eicosanoid species in normal and PanIN-bearing pancreata from 8-10 month old *Kras*^*G12D*^;*Ptf1a*^*Cre/+*^ (*KC*) mice (n = 5) and discovered significant upregulation of several eicosanoids, including TxB_2_, 6k-PGF1α, and PGD_2_; 5-HETE was found to be elevated but did not reach significance (Figure 1E)(DelGiorno et al., 2020a). To determine how eicosanoid species and levels change between pre-invasive PanIN and PDAC, we compared these data to the profiles we generated for *KPC* tumors. We observed a statistically significant gain in prostaglandins TxB_2_, 12-HHTrE, 6k-PGF1α, PGE_2_, and 8-HDoHE in tumors as compared to PanIN (Figure 1F). Interestingly, 12-HHTre appears to be tumor-specific. We next examined eicosanoid patterns across all the profiled samples using hierarchical clustering of select eicosanoids (Figure 1G). While normal samples clustered tightly and displayed low levels of selected eicosanoids, murine tumors displayed a large degree of heterogeneity. Notably, orthotopic tumors clustered by cell line, with those derived from FC-1199 displaying a trend towards increased eicosanoid production as compared to FC-1242 and FC-1245. While autochthonous tumors displayed an overall higher level of the selected eicosanoids, several samples clustered closer to orthotopic tumor samples. The five profiled PanIN samples displayed sample-to-sample heterogeneity but were characterized by intermediate levels of select eicosanoids (Figure 1G).

Altogether, these analyses identify classes of eicosanoids that are altered during pancreatic tumorigenesis. Specifically, several prostaglandins (PGE_2_, PGD_2_, TxB_2_, 12-HHTrE, 6k-PGF1α), HETEs, and HDoHEs are upregulated in murine tumor models in a relatively consistent manner, although there is heterogeneity within and between commonly used PDAC models (Figure 1H).

### Cell type-specific expression of eicosanoid synthases and receptors in murine PanIN

In our eicosanoid profiling studies, we identified significant changes in several eicosanoid species during pancreatic tumorigenesis (Figure 1G). As pancreas cellular composition changes with disease progression, the relative contribution of each of these various cell types (i.e., epithelium, inflammatory cells, fibroblasts) to eicosanoid production likely changes as well. To identify these changes and to infer the major cellular sources of key eicosanoids throughout disease progression, we examined available single cell RNA sequencing (scRNA-seq) datasets generated from murine models of PanIN and PDAC. We focused our analyses on eicosanoid synthases and receptors with well-defined biosynthetic pathways, including prostaglandins and lipoxygenases.

Schlesinger et al., recently generated an extensive scRNA-seq dataset comprised of 43,897 cells encompassing epithelial and stromal cell types from 9 *Kras*^*G12D*^;*Ptf1aCre*^*ERTM/+*^;*Rosa26*^*LSL-tdTomato/+*^ mice at various stages of disease progression (Schlesinger et al., 2020). To identify major cellular sources of eicosanoid synthesis and potential receptor interactions, we reanalyzed the Schlesinger dataset. As shown in Figure 2A, these sequence libraries were combined, and clusters were annotated by examining both classic cell type markers and previously published single-cell gene signatures identified in pancreas tissue (Figure S1B)(Elyada et al., 2019). As several time-points were included in this dataset, normal pancreas cells, pre-invasive cells (including tuft cells), and tumor cells are represented. We first examined expression of upstream synthases in the prostaglandin synthesis pathway. We detected expression of phospholipase *Pla2g4a* in epithelial cells - including pre-invasive cells and tuft cells - as well as in stromal cells such as fibroblasts (Figure 2B). The cyclooxygenases *Ptgs1* (COX1) and *Ptgs2* (COX2) are co-expressed in macrophages, fibroblasts, and tuft cells (as previously described) (Delgiorno et al., 2014). *Ptgs1* is additionally expressed in endothelial cells and several immune cell types (Figure 2B-C). We next examined expression of terminal prostaglandin synthases and identified macrophages and tuft cells to be major sources of the PGD_2_ synthase *Hpgds*, while ductal cells strongly express the less efficient PGD_2_ synthase Ptgds. All PGE_2_ synthases (*Ptges1-3*) and PGI_2_ synthase Ptgis are expressed in fibroblasts. The thromboxane (TxA_2_, TxB_2_) and 12-HHTre synthase *Tbxas1* is detected in macrophages, while 12-HHTre synthase *Cyp2s1* is expressed in PanIN and tumor cells (Figure 2B-C). In terms of lipoxygenase expression, 5-HETE requires co-expression of *ALOX5* and its activating protein *ALOX5ap*, which are both observed in macrophages, neutrophils, and tuft cells (Figure 2B)(Wang and Dubois, 2010). To confirm expression of ALOX5 in tuft cells, we conducted co-immunofluorescence for ALOX5, and COX1 and acetylated-α-tubulin, which label tuft cells in the epithelium. Consistent with studies conducted in the intestines (McGinty et al., 2020), we found that 84% (336/400) of tuft cells (n = 4 *KC* mice) express ALOX5. Conversely, 99.5% (398/400) of ALOX5-expressing cells in the epithelium are tuft cells, suggesting a role for pancreatic tuft cells in 5-HETE synthesis (Figure 2D).

**Figure 2.**
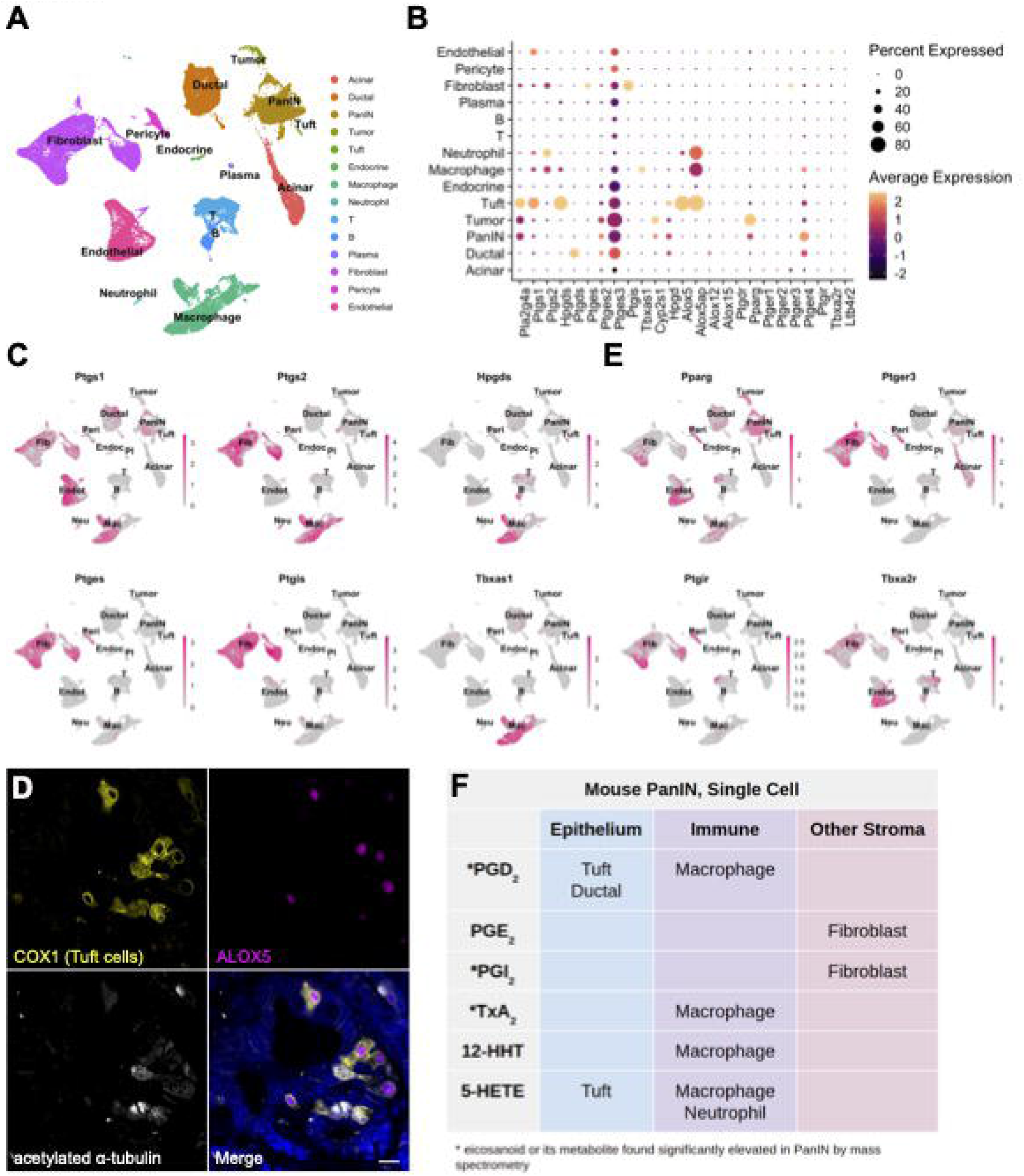
Cell type-specific expression of eicosanoid synthases and receptors in murine PanIN. **(A)** UMAP of a murine PanIN scRNA-seq dataset, generated from Schlesinger et al., annotated by cell type. **(B)** Dotplot of average and percent expression of select eicosanoid synthases and receptors in each cell type identified. **(C)** UMAPs depicting gene expression of select eicosanoid synthases. **(D)** Co-IF for COX1 (yellow), ALOX5 (magenta), and acetylated α-tubulin (white) in a *KC* pancreas. DAPI, blue. Scale bar, 10 μm. **(E)** UMAPs depicting gene expression of select eicosanoid receptors. Color intensity in (C) and (E) indicates the normalized gene expression level for a given gene in each cell. **(F)** Table summarizing inferred cellular sources of eicosanoids based on synthase expression patterns.

To determine which cell types might respond to relevant eicosanoids, we evaluated patterns of eicosanoid receptor expression. We identified widespread expression of receptor *Pparg* (PGD_2_ and PGE_2_ metabolites, 15-HETE) in normal and pre-invasive ductal cells as well as in several stromal populations. Expression of PGI_2_ receptor *Ptgir* (6k-PGF1α) is concentrated in fibroblasts and pericytes and the thromboxane receptor *Tbxa2r* (TxB_2_) is highly expressed in fibroblasts, pericytes, and endothelial cells. PGE_2_ receptors were also detected in our analysis, with *Ptger3* expression largely relegated to fibroblasts and *Ptger4* mainly expressed in normal and pre-invasive ductal cells, macrophages, T cells, and fibroblasts (Figure 2 B, E).

Collectively, our eicosanoid profiling and analysis of the Schlesinger et al. scRNA-seq dataset suggest potentially critical cellular sources of key eicosanoids in pre-invasive PanIN. For example, high levels of PGD_2_, PGI_2_ metabolites (6k-PGF1α), and TxA_2_ metabolites (TxB_2_) by mass spectrometry may be explained by the expression of requisite synthases in tuft cells and macrophages, fibroblasts, and macrophages, respectively (Figure 2B-C, 2F). Some of the cell types that may respond to these eicosanoids include fibroblasts (*Ptgir*, *Tbxa2r*), pericytes (*Ptgir*), and endothelial cells (*Tbxa2r*) (Figure 2B, E).

### Expression of eicosanoid synthases and receptors in murine models of PDAC

To examine eicosanoid synthase and receptor expression in murine PDAC, we reanalyzed a scRNA-seq dataset generated from *KPC* mice (Figure 3A)(Elyada et al., 2019). This dataset is comprised of 11,222 cells and libraries were combined and clusters annotated by expression of known cell type markers (Figure S1C). In contrast to the Schlesinger dataset, epithelial clusters in this dataset are largely comprised of tumor cells (Figure 3A). In terms of eicosanoid synthases, we identified expression of phospholipase *Pla2g4a* in tumor cells, fibroblasts, and myeloid cells, with minor expression in neutrophils (Figure 3B). Cyclooxygenases (*Ptgs1*, *Ptgs2*) are expressed primarily in EMT-like tumor cells, myeloid cells, neutrophils, and fibroblasts (Figure 3B). We identified the full complement of genes required to produce PGD_2_ (*Hpgds*), thromboxane pathway products TxB_2_ and 12-HHTre (*Tbxas1*), and 5-HETE (*ALOX5*, *ALOX5ap*), as well as *Alox15* (15-HETE, 14- and 17-HdoHE) in myeloid cells. Tumor cells express genes required to produce PGE_2_ (*Ptges*) and 12-HHTre (*Cyp2s1*). Although few fibroblasts were profiled in this dataset, enzymes required to produce PGE_2_ and PGI_2_ (*Ptgis*) were detected. Finally, neutrophils express enzymes required for 5-HETE synthesis (*ALOX5*, *ALOX5ap*) (Figure 3B-C).

**Figure 3.**
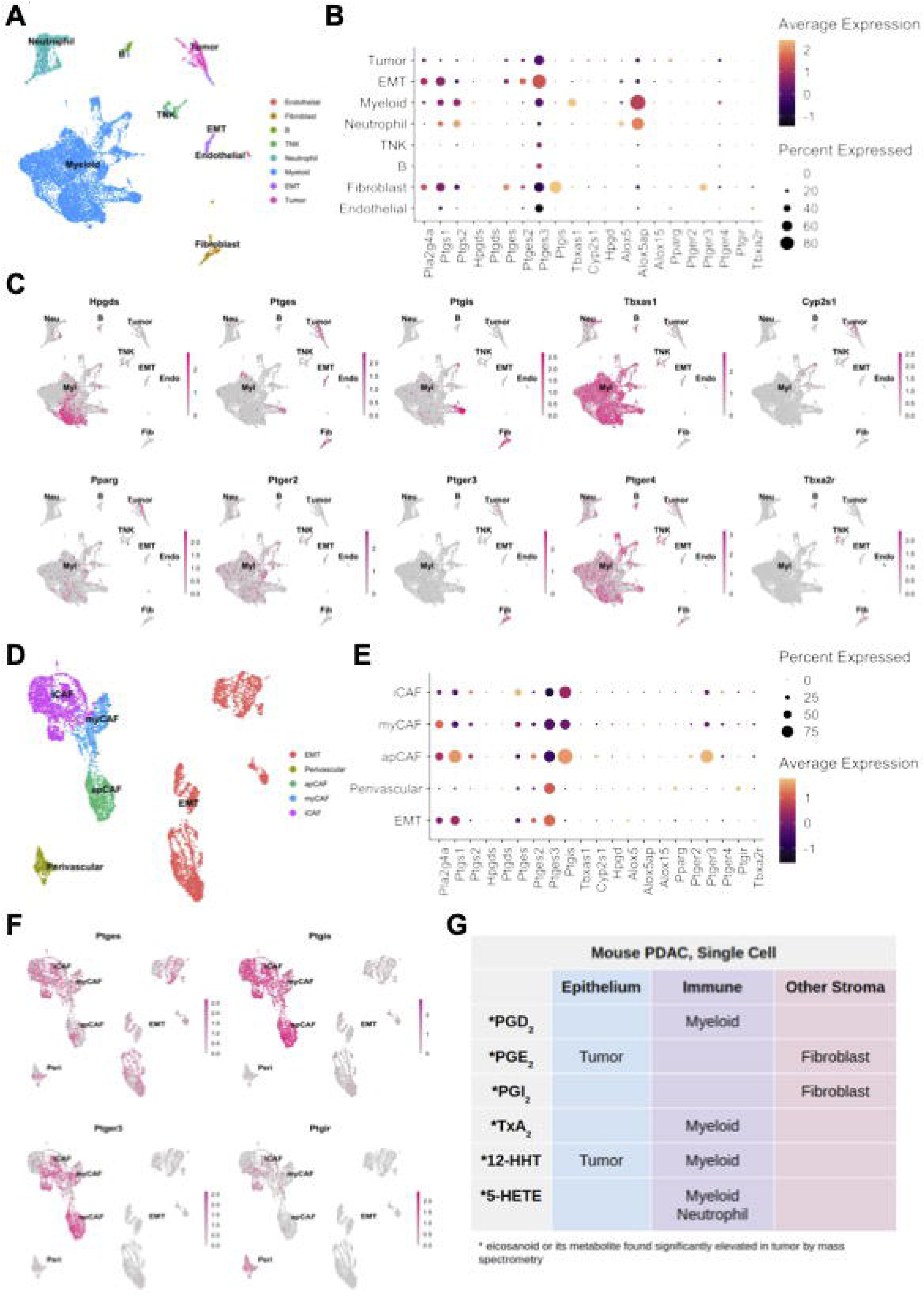
Cell type-specific expression of eicosanoid synthases and receptors in murine models of PDAC. **(A)** UMAP of a murine PDAC (*KPC*) scRNA-seq dataset derived from Elyada et al., annotated by cell type. EMT, epithelial to mesenchymal transition; TNK, T and natural killer cell. **(B)** Dotplot of average and percent expression of select eicosanoid synthases and receptors in each cell type. **(C)** UMAPs depicting gene expression of select eicosanoid synthases/receptors. **(D)** UMAP of a FACS-enriched CAF dataset derived from Elyada et al., annotated by cell type. iCAF, inflammatory cancer associated fibroblast; myCAF, myofibroblastic CAF; apCAF, antigen presenting CAF. **(E)** Dotplot of average and percent expression of select eicosanoid synthases and receptors in each cell type. **(F)** UMAPs depicting gene expression of select eicosanoid synthases/receptors. Color intensity in (C) and (F) indicates the normalized gene expression level for a given gene in each cell. **(G)** Table summarizing inferred cellular sources of eicosanoids based on patterns of synthase expression.

We next examined expression of known eicosanoid receptors. Pparg, which mediates the effects of several eicosanoids, is most highly expressed in tumor cells and myeloid cells (Wang and Dubois, 2010). PGE_2_ receptors are heterogeneously expressed in myeloid cells, tumor cells, neutrophils, and fibroblasts. While this dataset encompasses only a small number of endothelial cells, we could readily detect expression of both *Pparg* and thromboxane receptor *Tbxa2r*, consistent with the Schlesinger dataset (Figure 3B-C). To collectively validate our findings in the PanIN and PDAC datasets, we reanalyzed a third scRNA-seq dataset that encompasses normal pancreas, pre-invasive disease, and PDAC in multiple murine models (Figure S2)(Hosein et al., 2019). Several patterns found in the Schlesinger dataset were identified here as well, notably the presence of *Ptgds* and absence of *Ptges* in the epithelium in early stages of disease progression. Analyses of PDAC are largely in agreement with observations made from the Elyada et. al. dataset, including expression of PGE_2_ synthases in tumor cells, *Ptgis* in fibroblasts, and *Tbxas1* in macrophages/myeloid cells (Figure S2).

Cancer-associated fibroblasts (CAFs) have been established as important players in cancer formation and progression and several sub-sets have been identified in PDAC (Biffi et al., 2019; Elyada et al., 2019). To determine if eicosanoid synthases and/or receptors are differentially expressed in various CAF sub-types, we re-examined a second scRNA-seq dataset from Elyada et al., consisting of FACS-enriched fibroblasts. This dataset is composed of 8,438 cells generated from four tumor-bearing *KPC* mice and includes three CAF sub-types (iCAFs, myCAFs, apCAFs), perivascular cells, and EpCAM-negative EMT-like tumor cells, as annotated by the original authors (Figure 3D, S1D). We found eicosanoid synthase and receptor expression in this fibroblast-enriched dataset to be largely consistent with our previous analyses, showing that fibroblasts express *Pla2g4a*, *Ptgs1* (COX1), PGE_2_ synthases and *Ptgis*. Fibroblasts specifically express PGE_2_ receptor *Ptger3*, which is highest in apCAFs. Interestingly, the PGI_2_ (6k-PGF1α) receptor, *Ptgir*, is also enriched in perivascular cells suggesting a possible signaling loop between CAFs and perivascular cells (Figure 3E-F).

From these collective analyses, we infer cellular sources of pancreas disease-associated eicosanoids by examining patterns of eicosanoid synthase gene expression. The data suggests that, in PDAC, tumor cells are producing PGE_2_ and 12-HHTre, myeloid cells are producing PGD_2_, thromboxanes (TxA_2_, TxB_2_), 12-HHTre, and 5-HETE, and fibroblasts are generating PGE_2_ and PGI_2_ (6k-PGF1α)(Figure 3G).

To evaluate whether orthotopic tumor models recapitulate the patterns of eicosanoid synthase expression we identified in *KPC* mice, we generated tumors using the FC-1199 and FC-1245 PDAC cell lines. Tumors were collected and epithelial (EpCAM+CD45-) or inflammatory (EpCAM-CD45+) cells were isolated by FACS. Separately, RNA was collected from chunks of whole tumor or cell lines grown in 2-D. We then compared expression of eicosanoid synthases between these different models and tissue compartments by qRT-PCR and immunofluorescence. In tumors derived from both cell lines, we found eicosanoid synthases (*Ptgs1*, *Ptgs2*, *Hpgds*, *ALOX5*) to be more highly expressed in immune cells than in tumor cells, though expression is not exclusive (Figure 4A-C). PGD_2_ synthase *Ptgds* and PGE_2_ synthases **Ptges*1-3* are also expressed in tumor cells (Figure 4A-B). Fibroblasts are less abundant in these tumors than in autochthonous models, but we were able to identify expression of *Ptgis* in αSMA+ fibroblasts by immunofluorescence (Figure 4C). Notably, and in comparison, to autochthonous models and human disease (Figures 6-7), eicosanoid synthesis in orthotopic models is dominated by the stromal compartment rather than the tumor cells (Figure 4).

**Figure 4.**
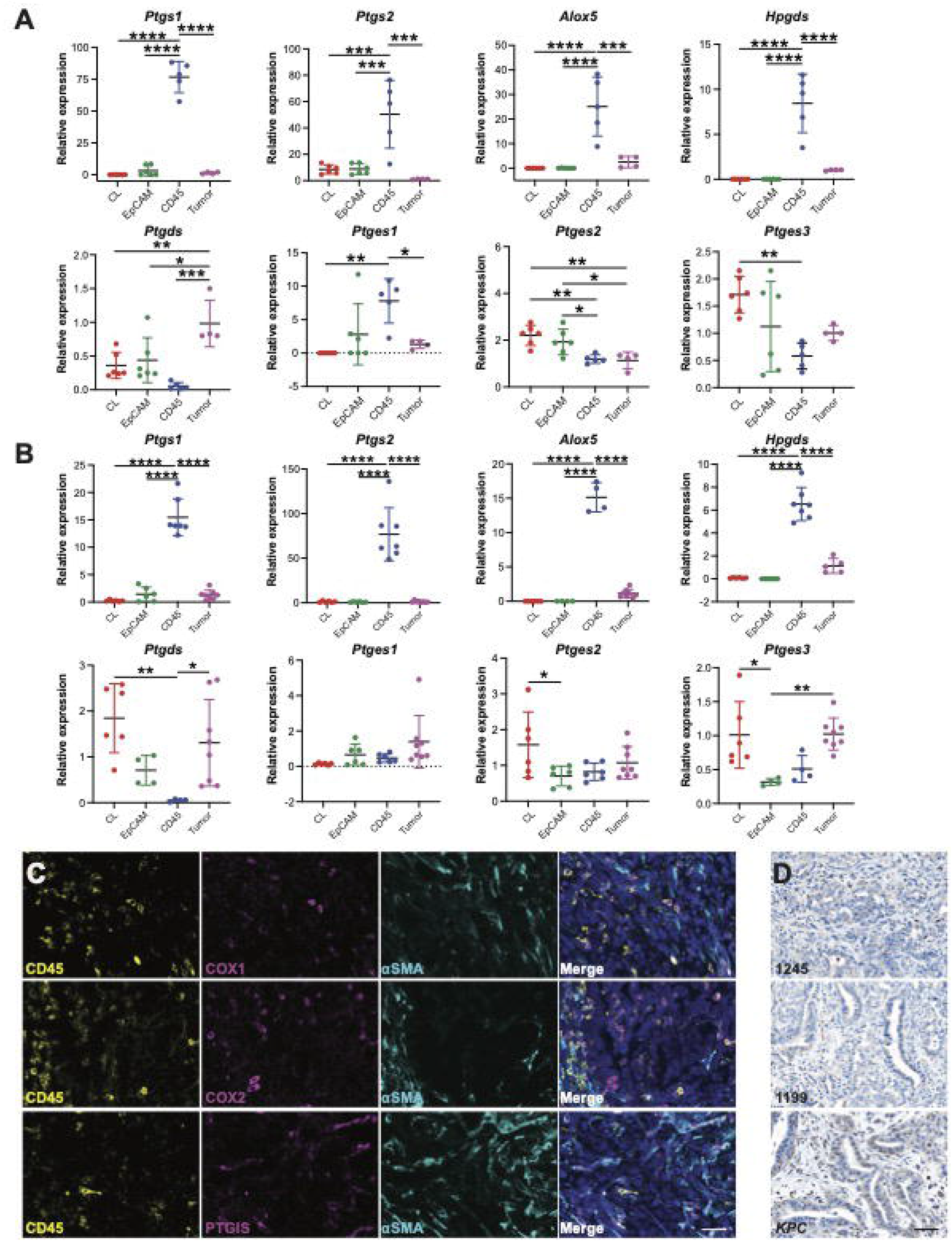
Cell type-specific expression of eicosanoid synthases in orthotopic models of PDAC. qRT-PCR for select eicosanoid synthases in **(A)** the FC-1245 or **(B)** the FC-1199 cell lines (CL), FACS collected EpCAM+ tumor or CD45+ immune cells, or whole orthotopic tumors. **(C)** Co-IF for COX1, COX2, or *Ptgis* (magenta) and CD45 (yellow), αSMA (cyan), and DAPI (blue) in orthotopic tumors generated from the FC-1245 cell line. **(D)** IHC for COX2 in orthotopic tumors generated from FC-1245 or FC-1199 or a *KPC* tumor. Scale bars, 50 μm. *, p < 0.05; **, p < 0.01; ***, p < 0.001; ****, p < 0.0001.

### Eicosanoid profiling of human PDAC

To determine if the eicosanoid profiles we generated from murine PDAC are representative of the human condition, we next conducted mass spectrometry on surgical specimens collected from 12 patients with PDAC, as well as 13 pancreas samples pathologically determined to be normal (Figure 5A). Eicosanoid profiling of human samples revealed large, significant upregulation of prostaglandins in the thromboxane pathway (TxB_2_, 12-HHTre) and the prostacyclin pathway (6k-PGF1α), consistent with observations made in mouse models (Figure 1). Though it did not reach significance, we did see a trend towards increased PGE_2_ levels in PDAC (p=0.059), in agreement with previous studies (Figure 5D)(Markosyan et al., 2019). Consistent with our observations in murine orthotopic tumor models, we observed a significant decrease in multiple linoleic acid metabolites in PDAC as compared to normal pancreas (Figure 5B-D). We also detected a significant decrease in free adrenic acid and docosahexaenoic acid (Figure 5D). Interestingly, and in agreement with our previous studies of HPGDS expression, we did not detect high levels of PGD_2_ (Figure 5D)(DelGiorno et al., 2020a). While this may be representative of human PDAC, we cannot exclude confounding factors, such as sample preparation. The lack of detection of additional eicosanoids found to be elevated in mouse models may be due, in part, to degradation.

**Figure 5.**
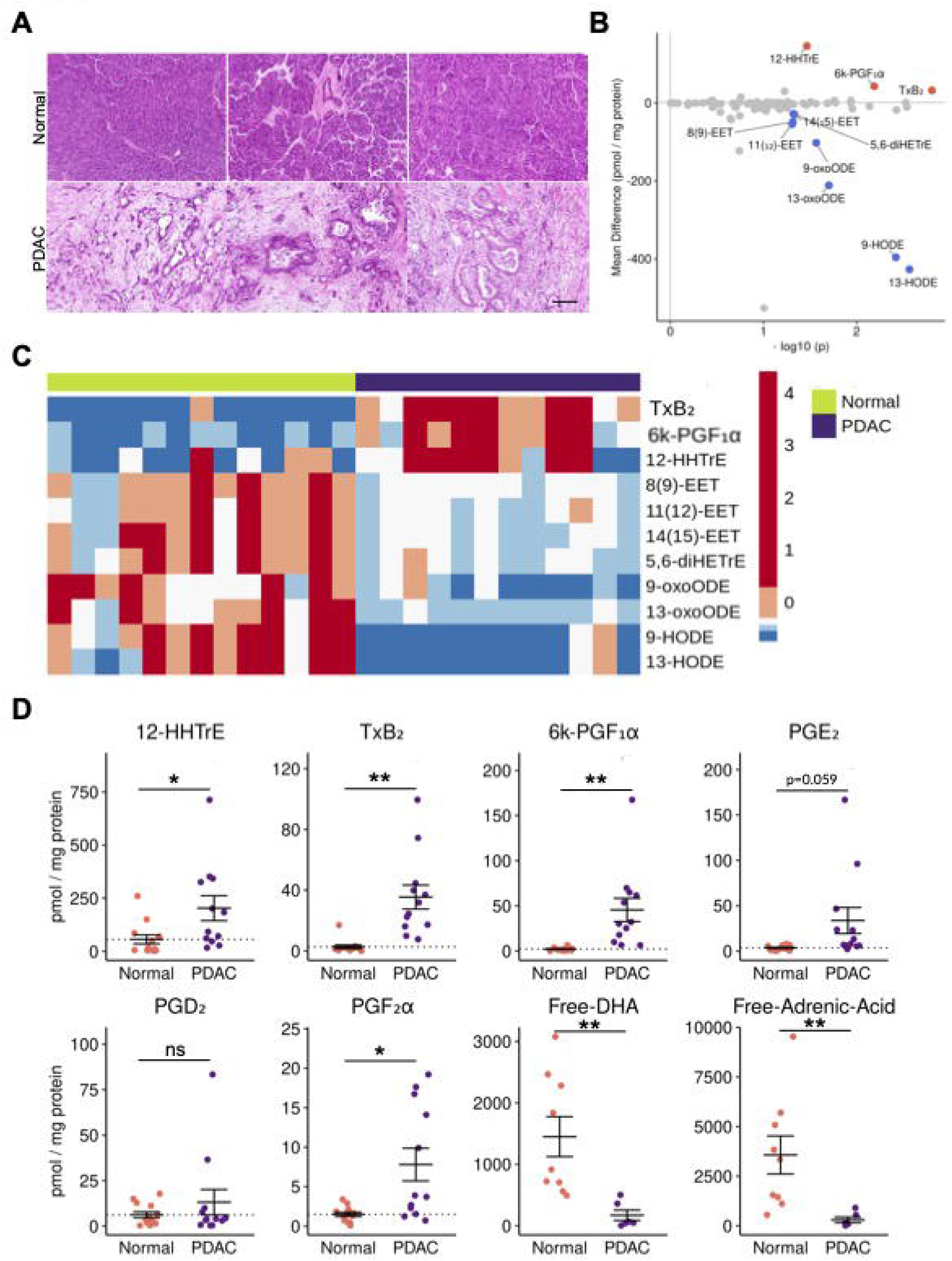
Eicosanoid profiling of human PDAC. **(A)** Representative H&E of normal human pancreata and PDAC samples. Scale bar, 100 m. **(B)** Comparison of normal pancreas and PDAC eicosanoid profiles (pmol/mg protein). Samples with a mean difference > +/− 20 pmol/mg protein and p < 0.05 are indicated in either red, up, or blue, down. **(C)** Heatmap of select eicosanoids from normal pancreata and PDAC. Values are z-score normalized by row and colors are assigned by quintile scaling. **(D)** Dotplots of select eicosanoids. Error bars, standard error of the mean. *, p < 0.05; **, p < 0.01; ***, p < 0.001; ****, p < 0.0001.

**Figure 6.**
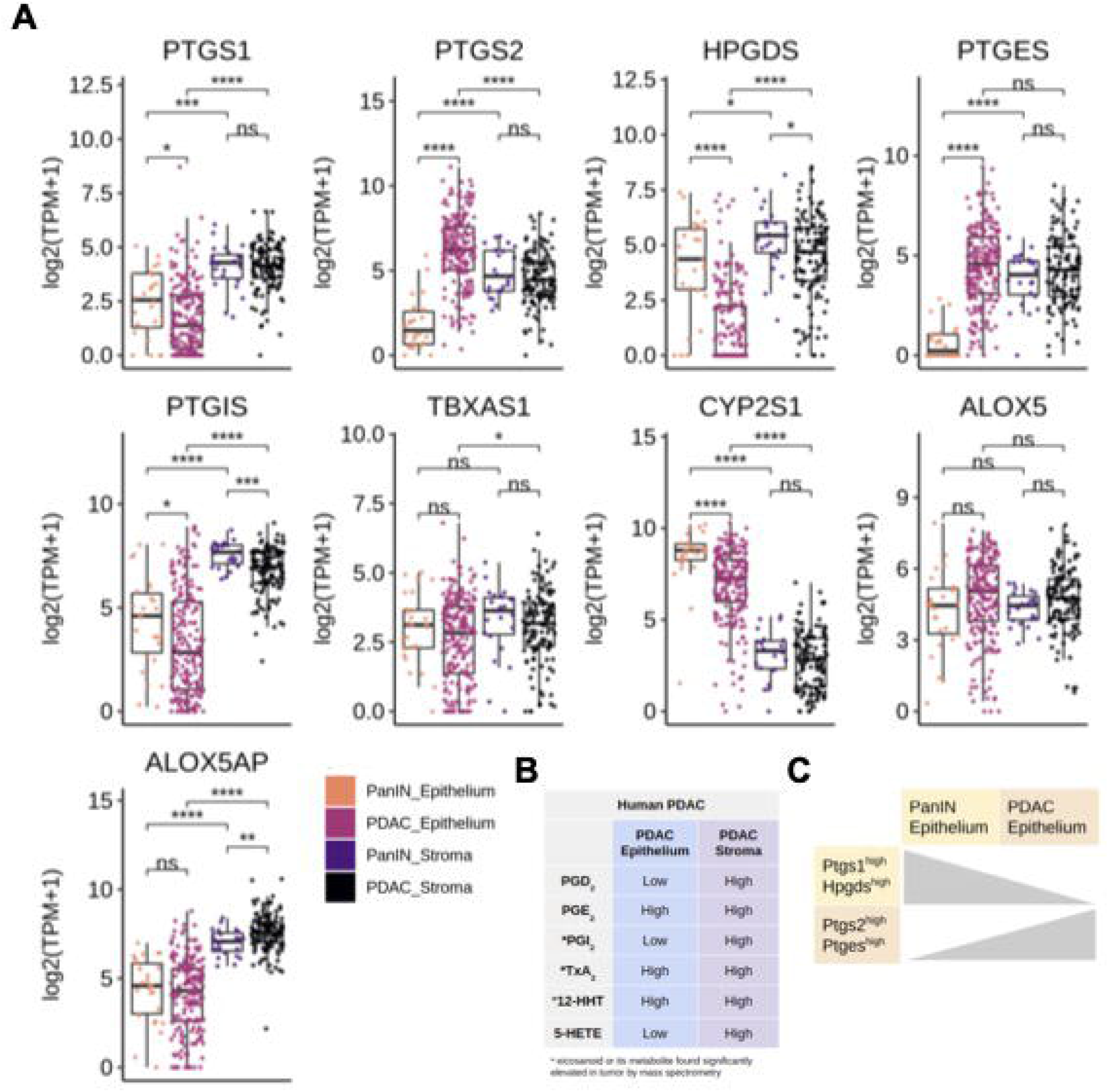
Eicosanoid synthase expression in human PanIN and PDAC. **(A)** Boxplots comparing expression (log2(TPM+1)) of microdissected stroma and epithelium from PanIN (n = 26) and PDAC (n = 197 epithelium, 124 stroma). *, p < 0.05; **, p < 0.01; ***, p < 0.001; ****, p < 0.0001. **(B)** Qualitative summary describing relative localization of terminal eicosanoid synthases in either the tumor epithelium or stroma. **(C)** Summary schematic describing a switch in prostaglandin synthase expression in the epithelium transitioning from PanIN to PDAC.

### Eicosanoid gene expression changes during the transition from pre-invasive PanIN to PDAC

As our eicosanoid profiling analysis of human PDAC identified several species that are also upregulated in mouse models, we next asked whether the cellular sources of these relevant eicosanoid synthases are the same and if they change during the PanIN to PDAC transition. To answer these questions, we examined a bulk RNA sequencing dataset of laser capture microdissected samples of human epithelium and stroma from 26 patients with PanIN and 197 patients with PDAC (124 matched stromal samples, generated as described in (Maurer et al., 2019)). First, we examined cyclooxygenase gene expression patterns. We found that while *PTGS1* (COX1) expression is higher in the stroma than the epithelium in both PanIN and PDAC, there is a significant decrease in gene expression in the epithelium from PanIN to PDAC, consistent with tuft cell loss (Figure 6A)(Delgiorno et al., 2014). Interestingly, there is a gain in *PTGS2* (COX2) expression in the epithelium in the transition to PDAC and levels are higher than in the stoma, consistent with a role for tumor cell-derived COX2 (Figure 6A).

Next, we examined expression patterns of terminal prostaglandin synthases. While both PGD_2_ synthase *HPGDS* and PGE_2_ synthase *PTGES* are expressed in the stroma of PanIN and PDAC, we found a large, significant decrease in *HPGDS* and a gain of *PTGES* expression in the PDAC epithelium (Figure 6A-C). These results suggest a possible switch between tumor suppressive PGD_2_ and pro-tumorigenic PGE_2_ during the transition from PanIN to PDAC.

Other features of human PDAC identified in this analysis are consistent with what we found in mouse models, namely higher expression of *PTGIS* in the stroma and *CYP2S1* in the epithelium (Figure 6A). Thromboxane synthase *TXBAS1* was detected at comparable levels in the epithelium of both PanIN and PDAC, but is significantly higher in the PDAC stroma, consistent with the increase in TxB_2_ levels we found between PanIN and PDAC in mouse models (Figure 6A). While *ALOX5* is expressed at comparable levels in all conditions, *ALOX5AP* is elevated only in stromal samples (Figure 6A). Taken together, these data show that the cell type-specific expression patterns of eicosanoid synthases in human PanIN and PDAC largely agree with that in mouse models, though tumor cells may play a larger role in human disease (Figure 6B, S3). Over the course of pancreatic tumorigenesis, we observed a shift in the epithelium from *PTGS1*/*HPGDS* to *PTGS2*/*PTGES* expression (Figure 6C). This is consistent with our observation that *PTGS1*+*HPGDS*+ tuft cells are lost in the transition from pre-invasive disease to PDAC (Delgiorno et al., 2014).

### Cell type-specific expression of eicosanoid synthases and receptors in human PDAC

To identify the specific epithelial and stromal cell types that express relevant eicosanoid synthases and receptors in human PDAC, we next examined a scRNA-seq dataset of 11 normal pancreata and 24 PDAC samples including 57,530 cells (Peng et al., 2019). We first subsetted the tumor samples (41,986 cells) and identified cell types by markers described in Peng et al. (Figure 7A). Interestingly, we identified a large degree of heterogeneity in synthase expression among malignant tumor cells, possibly reflecting the inter-tumor heterogeneity of patients included in this study (Figure 7B-C). Within the tumor population, we identified expression of phospholipase enzymes *PLA2G2A* and *PLA2G4A* and *PTGS2* (COX2), but not *PTGS1* (COX1) (Fig 7B-C). Tumor cells express synthases *TBXAS1* (TXA_2_, TXB_2_, 12-HHTre) and *CYP2S1* (12-HHTre) and some subclusters are enriched for PGE_2_ synthase *PTGES* or prostacyclin synthase *PTGIS* (Figure 7B-C). A small subset of 49 cells that cluster with ductal cells are tuft cells and express both *PTGS1* and *HPGDS*, as previously described (Figure S4A)(Delgiorno et al., 2014). In terms of the stroma, we identified expression of *PTGS1*, *TBXAS1*, and *CYP2S1* enriched in macrophages. A subset of these cells also weakly express *HPGDS*. PDAC CAFs express *PTGDS* (PGD_2_), *PTGES* (PGE_2_), and *PTGIS* (PGI_2_) (Figure 7B-C). *ALOX5*/*ALOX5AP* (5-HETE) co-expression is largely detected in myeloid cells, but also in B cells. Consistent with the Maurer dataset, *ALOX15B* (15-HETE, 17-HdoHE) is expressed in myeloid cells and fibroblasts (Figure 7B, S3)(Maurer et al., 2019). To confirm these data, we interrogated a second scRNA-seq dataset of human PDAC containing 8000 cells and 10 patients (Lin et al., 2020). As shown in Figure S4B, we found that eicosanoid synthase expression patterns in these two datasets are largely in agreement (Peng et al., 2019).

**Figure 7.**
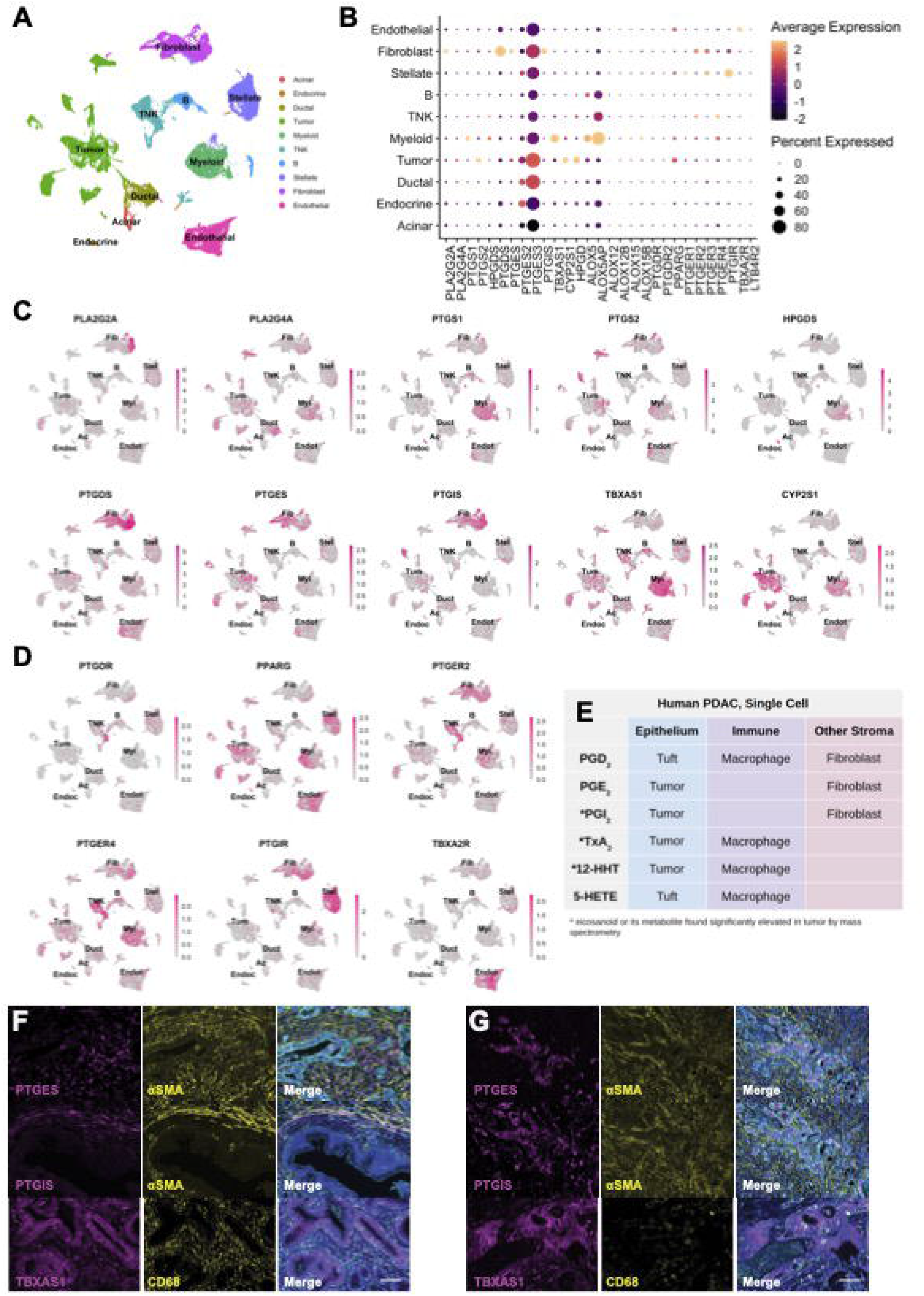
Cell type-specific expression of eicosanoid synthases and receptors in human PDAC. **(A)** UMAP of a human PDAC scRNA-seq dataset generated by Peng et al., subsetted to exclude adjacent normal samples and annotated by cell type. TNK, T and natural killer cells. **(B)** Dotplot of average and percent expression of select eicosanoid synthases/receptors in each cell type. UMAPs of either eicosanoid **(C)** synthase or **(D)** receptor gene expression. Color intensity indicates the normalized gene expression level for a given gene in each cell. **(E)** Table summarizing inferred cellular sources of eicosanoids based on patterns of synthase expression. **(F)** Co-immunofluorescence for relevant eicosanoid synthases (magenta), stromal markers (αSMA, fibroblasts; CD68, macrophages, yellow), and γactin (blue) highlighting stromal or **(G)** tumor expression. Scale bar, 50 μm.

To infer what cell types might respond to eicosanoids in the PDAC microenvironment, we next examined expression of relevant receptors. *PPARG* (PGD_2_ and PGE_2_ metabolites, 15-HETE) expression is widespread and can be found in tumor cells, myeloid cells, stellate cells, and endothelial cells, with minor expression in several other cell types. *PTGER2* and *PTGER4* share similar patterns of expression, highest in myeloid cells, T cells, and fibroblasts, while *PTGER3* is specifically expressed in stellate cells and fibroblasts. PGD_2_ receptor *PTGDR* is expressed in T cells, fibroblasts, and stellate cells. As seen in murine models, *PTGIR* (PGI_2_, 6k-PGF1α) is highly expressed in stellate cells and fibroblasts and *TBXA2R* (TxB_2_, TxA_2_) is most highly expressed in endothelial cells (Figure 7D).

To reconcile inferences made from the RNA-seq studies (Figure 6B, 7E) with eicosanoids detected by mass spectrometry (Figure 5), we conducted immunostaining for PTGIS, PTGES, and TBXAS1 in several tissue samples spanning normal pancreas, PanIN, and PDAC. Consistent with sequencing studies, we identified expression of PTGIS and PTGES in both αSMA+ fibroblasts and in tumor cells. TBXAS1 is expressed in both CD68+ macrophages and the tumor epithelium (Figure 7F-G).

To take a more quantitative approach, we next conducted immunohistochemistry (IHC) and scored expression of these synthases in multiple tissue compartments of samples from 22 treatment naïve PDAC patients. By sampling multiple surgical samples from the same patients, our analysis spanned normal tissue, metaplasia, high and low grade PanIN, well and poorly differentiated PDAC, and myxoid or compact stroma totaling 1176 regions of interest (Figure 8A, File S2). We note several observations from these analyses, including a significant increase in PTGES expression in the epithelium with disease progression, as predicted in Figure 6, highest in poorly differentiated PDAC. We also identified a significant increase in PTGIS in the stroma and in TBXAS1 in the epithelium of high grade PanIN, as compared to normal tissue, (Figure 8A-B). Using the same data, Spearman Rho correlation analyses between all the datapoint classes identify significant positive and inverse correlations mostly in interrogated normal duct areas (Figure S5A). Notably, PTGIS, PTGES, and TBXAS1 show positively correlated expression in both epithelial and stromal components (p=0.005-0.003×10^−5^), while PTGIS and TBXAS1 have an inverse correlation in normal duct epithelium (p=0.004). Furthermore, the stroma underlying poorly differentiated PDAC showed an inverse relationship between PTGIS and PTGES (p=0.032).

**Figure 8.**
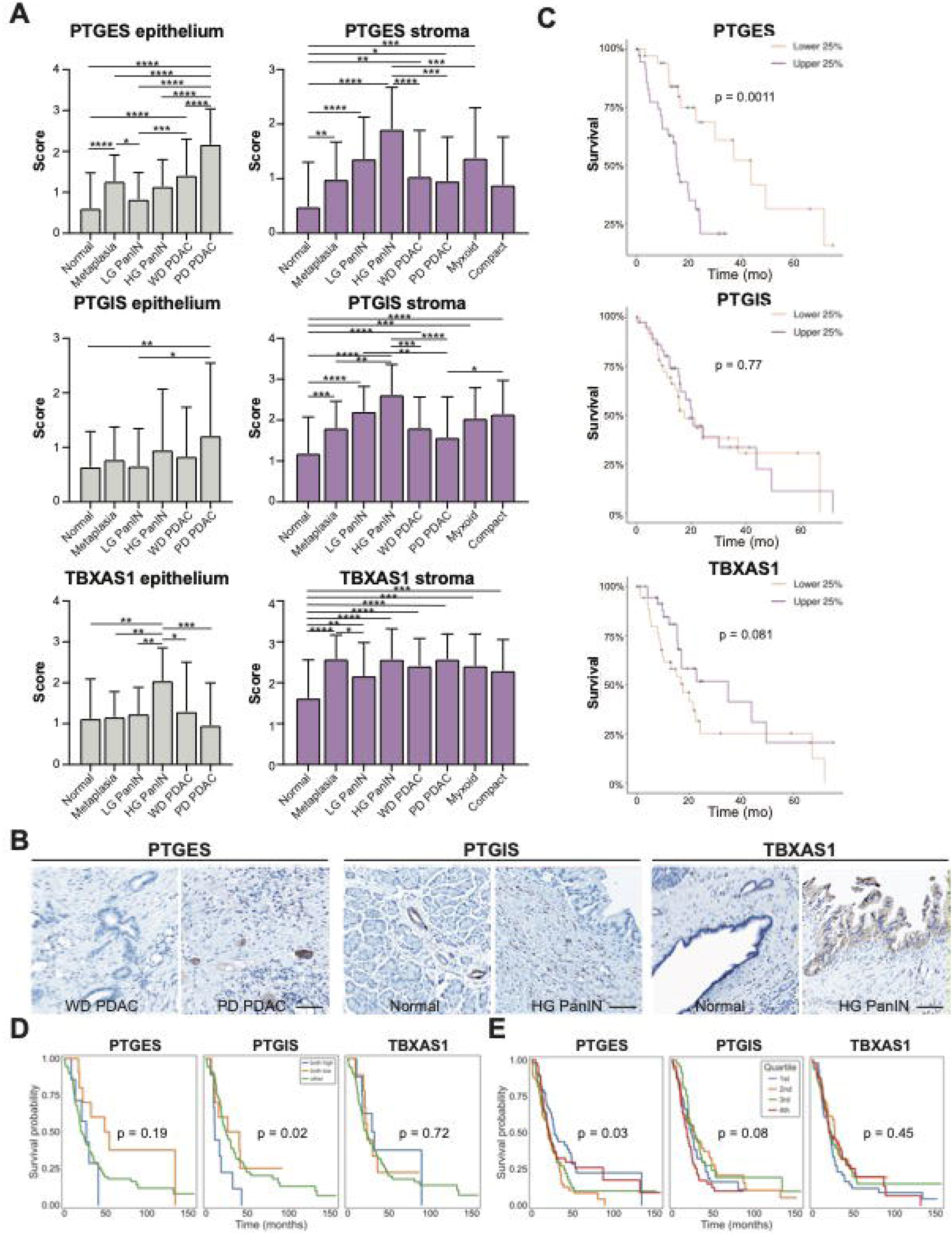
Localization and survival benefit for select eicosanoid synthases in human PDAC. **(A)** Quantification of scored IHC for PTGES, PTGIS, or TBXAS1 from 22 patients encompassing normal, PanIN, and PDAC-associated stroma and epithelium as well as both myxoid and compact fibrosis. **(B)** Representative IHC images. Scale bar, 100 m. Image diameter, 500 m. **(C)** Survival curves for total PTGES, PTGIS, or TBXAS1 expression generated from the TCGA database, representing 150 PDAC patients. The top and bottom 38 patients are shown. Survival curves for **(D)** total or **(E)** epithelium-specific PTGES, PTGIS, or TBXAS1 expression generated from the Maurer et al., dataset representing 197 epithelial samples and 124 stromal samples from PDAC patients.

Finally, to determine if synthase expression correlates to patient survival, we interrogated the TCGA database (150 PDAC patients) as well as the laser-captured RNA-seq dataset (Maurer et al., manuscript in preparation, 197 PDAC epithelium, 124 PDAC stroma samples) described in Figure 6. As shown in Figure 8, we identified a survival advantage for patients with low PTGES expression in both datasets, and for patients with low PTGIS expression in the Maurer dataset (Figure 8C-D). Interestingly, it is tumor cell, and not stromal, expression of PTGES that is associated with a survival advantage in the Maurer dataset (Figure 8D-E, S5A). Collectively, these data suggest that tumor cell expression of eicosanoid synthases is associated with pathogenesis and reduced patient survival.

## Discussion

In this study, we conducted eicosanoid profiling on normal pancreata and PDAC in mouse models of pancreatic tumorigenesis and patient samples, interrogated published RNA-seq datasets to generate predictions as to the cellular origin of eicosanoid species, and validated key findings at the protein level. The data collectively suggest a previously undescribed role for prostacyclin and thromboxane signaling in pancreatic tumorigenesis. Transcriptomic data from both human and mouse samples suggest an “eicosanoid switch” in the epithelium, from PGD_2_-producing enzymes in PanIN to PGE_2_-producing enzymes in PDAC. Our analysis of the human condition suggests that this gain in synthase expression in tumor cells is associated with reduced survival. In mouse models, the stroma appears to be the dominant source of eicosanoid synthesis, a potentially critical difference between mouse models and human disease. Specifically, in mouse models of PDAC, we have found significant heterogeneity and diversity of secreted eicosanoid species – including prostaglandins, HETEs, and HdoHEs - across model types and gene-specific cre drivers. By integrating RNA transcript expression, eicosanoid levels, and whenever possible, synthase protein expression, we have catalogued the patterns of eicosanoid production and receptor expression during pancreatic disease and tumorigenesis (summarized in Figure 9). These analyses provide a valuable resource for further functional investigation.

**Figure 9.**
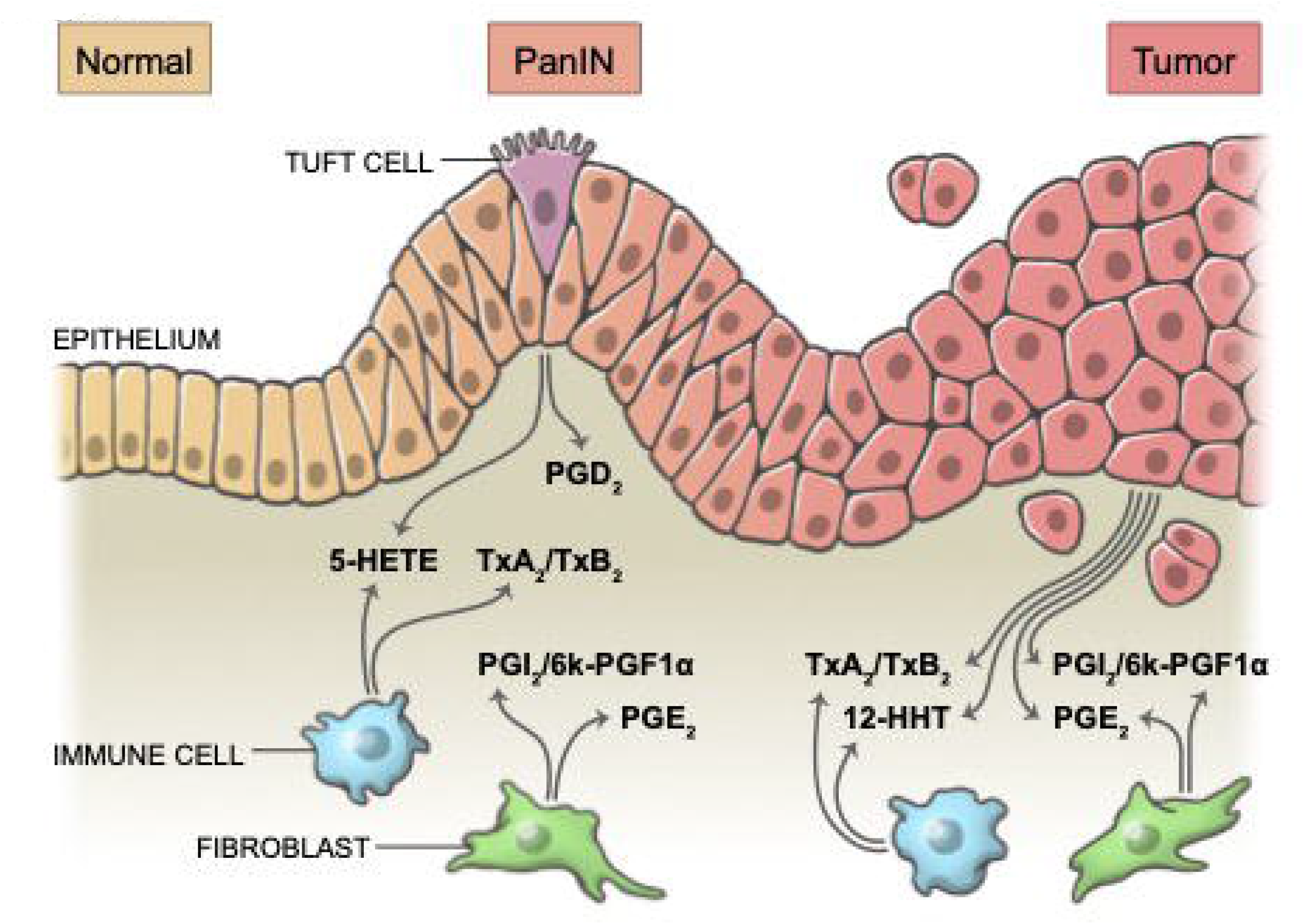
Summary of select eicosanoids and the cell types responsible for their synthesis throughout pancreatic tumorigenesis. Mass spectrometry identified minimal levels of eicosanoid species in the normal pancreas, however many species are significantly elevated as the pancreas undergoes tumorigenesis. We’ve previously shown that PanIN tuft cells generate PGD_2_; gene expression and immunostaining suggest that tuft cells also secrete *ALOX5*-derived 5-HETE. PanIN and PDAC-associated fibroblasts likely secrete both PGE_2_ and PGI_2_/6k-PGF1α and macrophages are a major source of TBXAS1-derived TxA_2_/TxB_2_. As disease progresses, the epithelium increases expression of PTGES (PGE_2_), PTGIS (PGI_2_/6k-PGF1α), and TBXAS1 (TxA_2_/TxB_2_ and 12-HHT).

We note certain limitations in the conclusions drawn from the experimental and analytical techniques we have used. Although eicosanoid profiling by mass spectrometry informed us of targets highly upregulated in PDAC, the detection of these species may have been impacted by their half-lives. For example, while it can be hypothesized that the relative lack of diversity in eicosanoid species in human PDAC may be a result of years-long immune reprogramming to an immunosuppressed state, in contrast to the months-long time course of tumor progression in mouse models, an equally viable hypothesis is that this lack of eicosanoid diversity is merely a technical artifact arising from sample processing post-surgery. This differs from the relatively quick preservation of mouse tumor samples. In support of the latter hypothesis is the fact that the three significantly upregulated eicosanoids – or their metabolites – in human tumors have relatively long half-lives. However, the trend towards higher PGE_2_ levels, which has a half-life of only 2.5-5 minutes may serve as a positive control (Bygdeman, 2003). These hypotheses may be resolved by accurate *in situ* studies aimed at identifying synthase functionality.

RNA sequencing based analyses also present key limitations. RNA expression does not always correlate with protein expression or activity, the latter of which is largely dependent on post-transcriptional factors, post-translational factors, enzyme localization and substrate availability, and signaling cues. Secondly, the scRNA-seq datasets we used to localize eicosanoid synthase/receptor expression are unsuitable to study platelets, whose small size and lack of a nucleus excluded them from collection. Platelets synthesize eicosanoids (PGE_2_, PGD_2_, 11-HETE, 15-HETE, and others) in various contexts, such as wound healing, and tumor infiltrating platelets have been described in pancreatic neuroendocrine tumors (Crescente et al., 2019; Xu et al., 2019). Activated platelets could represent a major source of eicosanoid diversity uncharacterized by our analyses.

Despite these limitations, our analyses identify eicosanoids that warrant further investigation in the context of pancreatic tumorigenesis. Our multifaceted approach enabled us to make novel predictions regarding patterns of eicosanoid synthesis in PanIN and PDAC. We have confirmed these predictions at the synthase RNA, synthase protein, and eicosanoid levels. Notably, many patterns of eicosanoid synthase expression identified in this study are consistent with previous studies in PDAC or in other organ systems. In addition to the numerous studies pointing to a role for tumor-cell derived PGE_2_, a recent study describes the ability of intestinal fibroblasts to secrete PGE_2_, which accelerates tumorigenesis (Nakanishi and Rosenberg, 2013; Roulis et al., 2020). While prostacyclin synthesis has been examined in the context of skin fibroblasts and dental cyst fibroblasts, our work suggests PDAC CAFs may be producing prostacyclin as well (Baenziger et al., 1979; Harvey et al., 1984). Finally, macrophages are well known sources of prostaglandins in a variety of organ systems, and have been shown to secrete PGD_2_, PGE_2_, and thromboxanes (Stachowska et al., 2009; Tripp et al., 1988; Virtue et al., 2015). Collectively, our work localizes eicosanoid synthases to specific cell types in PDAC and underscores the need to determine the function of these species in pancreatic cancer development and progression. Understanding the role of various eicosanoids in PDAC may identify pathways to co-opt or novel targets for treatment.

## Author contributions

Conceptualization: K.E.D. Formal analysis: V.B.G., N.J., V.Q.T., Z.M., H.C.M., W.L., P.K.S., K.E.D. Funding acquisition: H.H., T.H., P.K.S., S.M.K., K.P.O., G.M.W., K.E.D. Investigation: V.B.G., N.J., V.Q.T., R.F.N., N.K.L., C.H.M., S.Z., W.L., Y.S., K.E.D. Project administration: K.E.D. Resources: H.C.M., W.L., H.H., Y.S., T.H., K.P.O. Supervision: H.H., T.H., M.C.B.T., P.K.S., S.M.K., K.P.O., G.M.W., K.E.D. Visualization: V.B.G., K.E.D. Writing: V.B.G., G.M.W., K.E.D.

## Data and materials availability

All relevant data are included in this manuscript.

## Acknowledgements

The authors thank Taylor Culpepper for technical assistance and Amy Cao for artistic rendering and illustrator services. Tissue samples were provided by the NCI Cooperative Human Tissue Network (CHTN). Other investigators may have received specimens from the same tissue specimens. N.K.L. was supported by the Salk Institute Cancer Training Grant T32 CA009370, a Hope Funds for Cancer Research Postdoctoral Fellowship (HFCR-20-03-03), and a Sky Foundation Seed Grant. S.Z. was supported by NIH/NCI K00CA222741. The Flow Cytometry Core Facility at the Salk Institute is funded by NIH/NCI P30 CA014195 and Shared Instrumentation Grant S10-OD023. P.K.S. is supported by funding from the NIH/NCI (R01CA210439, R01CA163649, R01CA216853, NCI) and the Fred & Pamela Buffett Cancer Center Support Grant (P30CA036727). Work in the Olive laboratory is supported by the Herbert Irving Comprehensive Cancer Center Support Grant (P30CA013696) and the Pancreas Center of New York Presbyterian Hospital. M.C.B.T is supported by a Vanderbilt Digestive Disease Research Center Pilot and Feasibility Grant (P30 DK058404) and a Nikki Mitchell Foundation Pancreas Club Seed Grant. The Hunter laboratory was supported by NIH/NCI CA082683 and a Lustgarten Foundation Award (#552873). T.H. is a Frank and Else Schilling American Cancer Society Professor and the Renato Dulbecco Chair in Cancer Research. The Kaech laboratory was supported by NIH/NCI 5R01 CA240909-02. Work in the Wahl laboratory is supported, in part, by the Salk Institute Cancer Center Core Grant (CA014195), NIH-NCI R35 CA197687, NIH-NCI T32 CA009370, NIH/NCI 5R01 CA240909-02, the Isacoff Gastrointestinal Research Foundation, the Freeberg Foundation, and the Leona M. and the Harry B. Helmsley Charitable Trust (2012-PG-MED002). The DelGiorno laboratory is supported by NIH/NCI 5R01 CA240909-02, the Vanderbilt-Ingram Cancer Center Support Grant (P30 CA068485), a Vanderbilt Digestive Disease Research Center Pilot and Feasibility Grant (P30 058404), an American Gastroenterological Association Research Scholar Award (AGA2021-13-02), and Linda’s Hope (Nashville, TN). The DelGiorno laboratory receives funding from Cumberland Pharmaceuticals. The authors have no additional financial interests.

---

**Figure S1.**
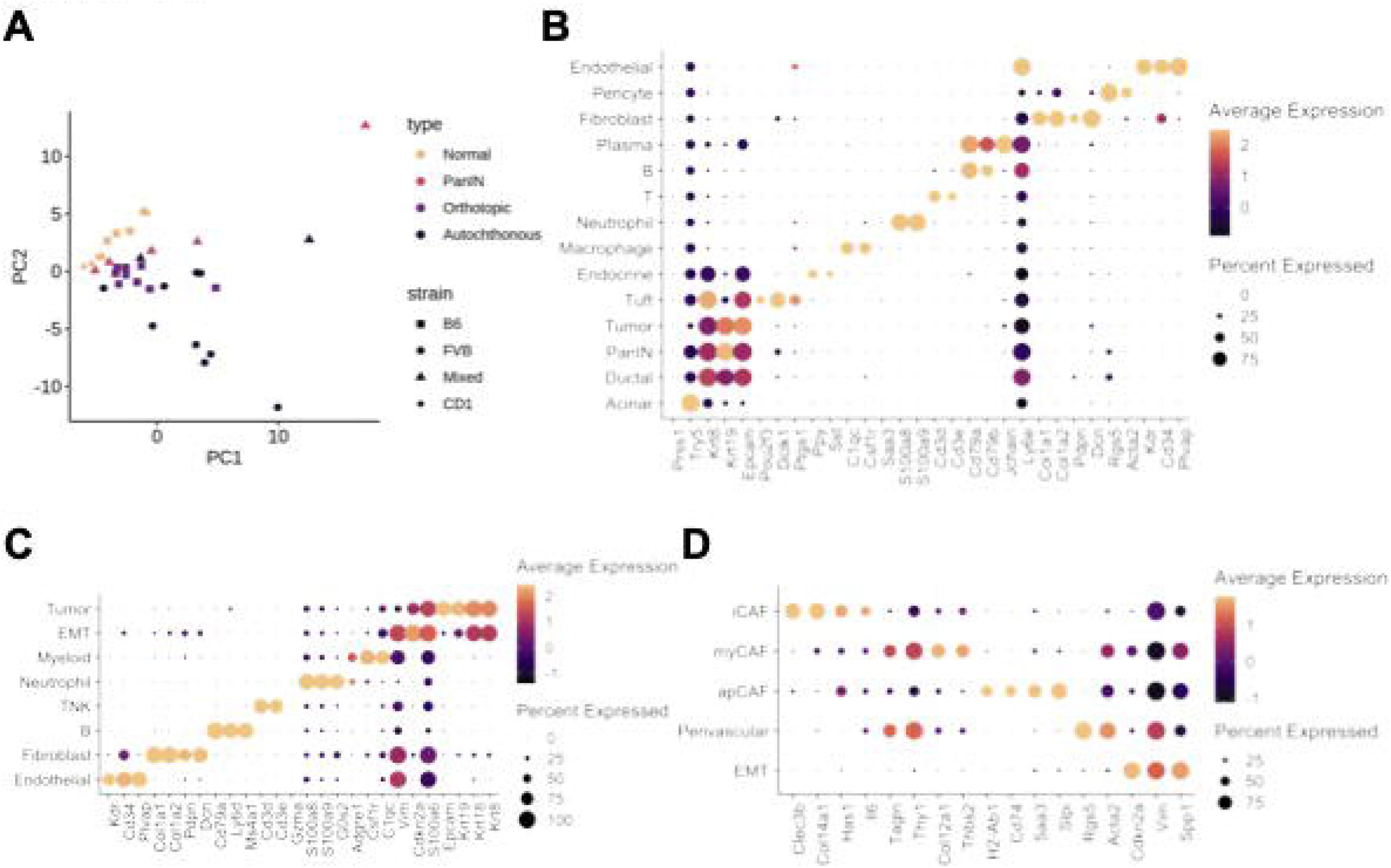
Eicosanoid signatures and cell type-specific markers in murine models of pancreatic tumorigenesis. **(A)** PCA plot comparing eicosanoid signatures of normal pancreata, PanIN-bearing pancreata, orthotopic PDAC tumors, or autochthonous PDAC tumors. **(B)** Dotplot of average and percent gene expression for cell type markers in a scRNA-seq dataset derived from Schlesinger et al. **(C)** Dotplot of average and percent gene expression for cell type markers in the entire unenriched dataset and (D) the fibroblast-enriched dataset generated from Elyada et al. EMT, epithelial to mesenchymal transition; TNK, T and natural killer cells; iCAF, inflammatory cancer-associated fibroblast; myCAF, myofibroblastic CAF; apCAF, antigen-presenting CAF.

**Figure S2.**
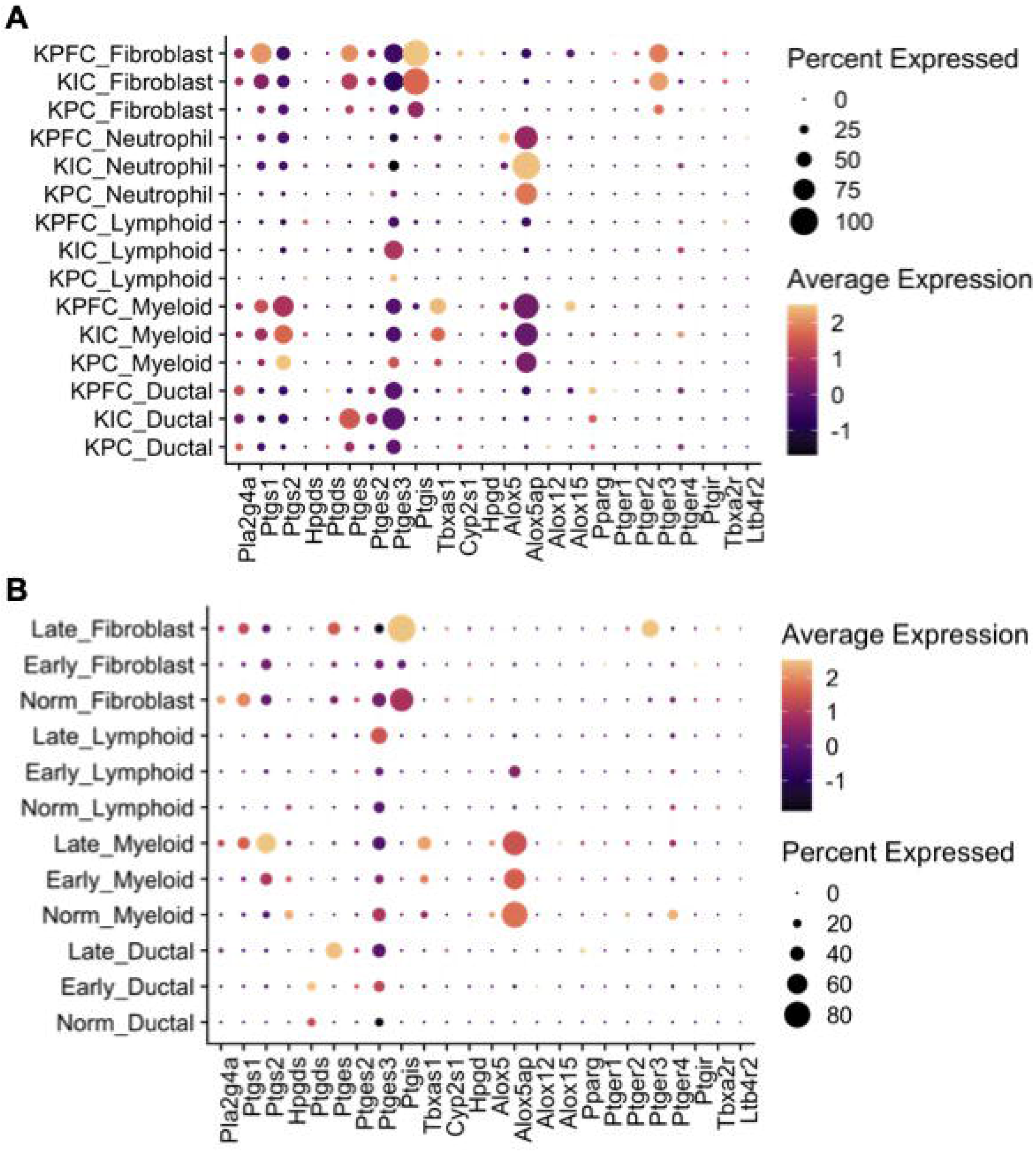
Eicosanoid pathway gene expression in multiple murine models of PDAC. Dotplots of eicosanoid synthase and receptor gene expression in normal pancreas and tumor epithelium organized by **(A)** mouse model or **(B)** stage of disease progression from the datasets described in Hosein et. al. *KPC*, *LSL-Kras*^*G12D*^;*Trp53*^*R172H*^;*Ptf1a*^*Cre/+*^; KIC, *LSL-Kras*^*G12D*^;*Ink4a*^*fl/fl*^;*Ptf1a*^*Cre/+*^; KPFC, *LSL-Kras*^*G12D*^;*Trp53*^*fl/fl*^; *Pdx1*^*Cre/+*^. Norm, normal; Early, early lesions; Late, PDAC.

**Figure S3.**
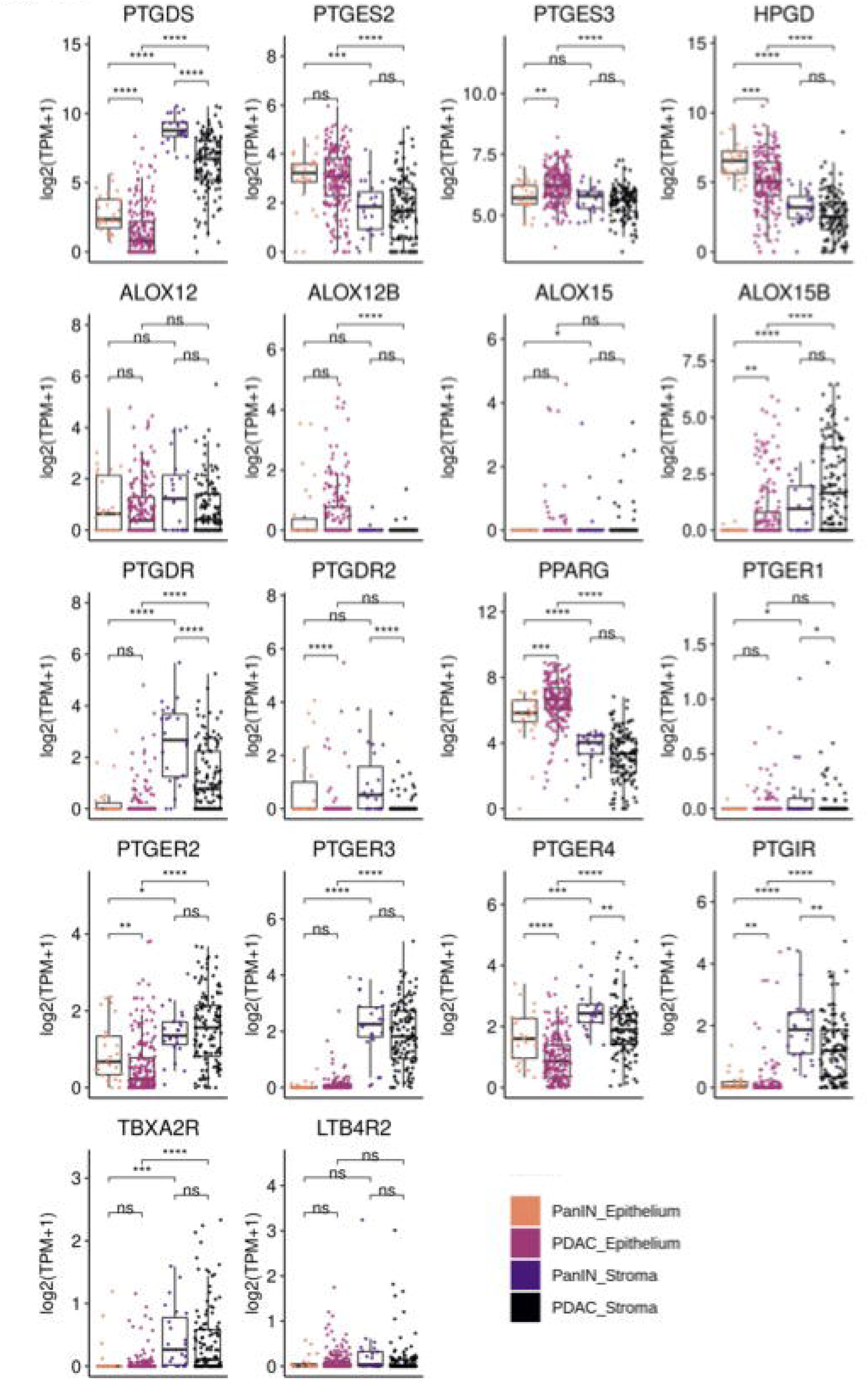
Eicosanoid pathway gene expression in the epithelium or stroma of human pre-invasive disease and PDAC. Boxplots comparing gene expression of eicosanoid synthases (log2(TPM+1)) in microdissected stroma and epithelium from PanIN or PDAC. *, p < 0.05; **, p < 0.01; ***, p < 0.001; ****, p < 0.0001.

**Figure S4.**
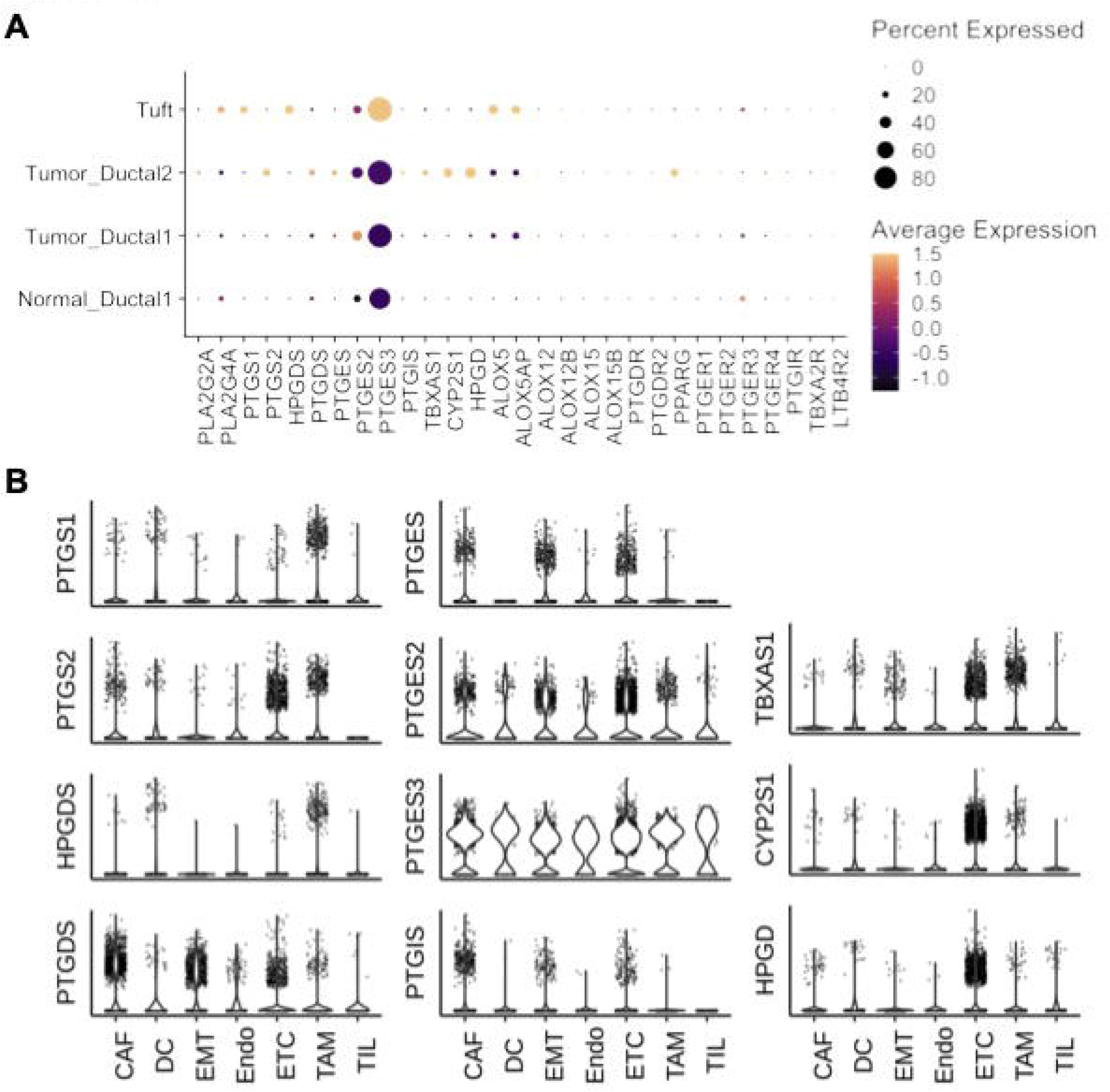
Eicosanoid synthase expression in human PDAC. **(A)** Heatmap of eicosanoid synthase and receptor gene expression in normal pancreas and tumor epithelium, from the dataset described in Peng et al. **(B)** Violin plots of eicosanoid synthase expression from a human PDAC scRNA-seq dataset described in Lin et al. CAF, cancer-associate fibroblast; DC, dendritic cells; EMT, epithelial to mesenchymal transition; Endo, endothelial; ETC, epithelial tumor cells; TAM, tumor-associated macrophages; TIL, tumor-associated lymphocytes.

**Figure S5.**
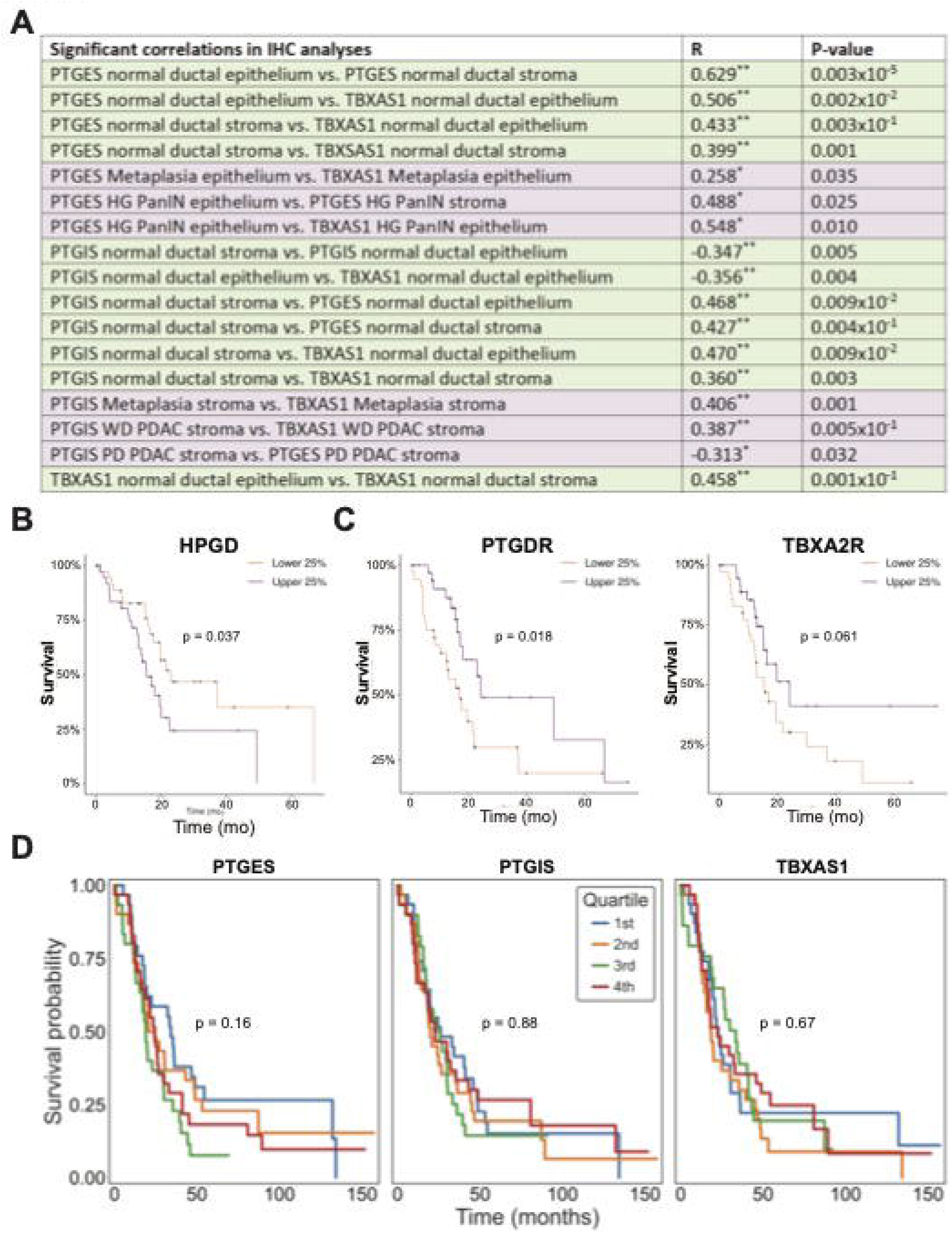
Immunohistochemical analysis of eicosanoid sythases in PDAC samples and patient survival associated with expression of select eicosanoid pathway genes. **(A)** Table of significant correlations identified in the IHC analysis of PTGES, PTGIS, and TBXAS1 expression. Green, normal tissue; purple, diseased tissue. HG, high grade; WD, well differentiated; PD, poorly differentiated. **(B)** Survival curve for HPGD expression generated from the TCGA database representing 150 PDAC patients. The top and bottom 38 are shown. **(C)** Survival curves for eicosanoid receptors *PTGDR* or *TBXA2R* generated from the TCGA database. **(D)** Survival curves for stroma-specific expression of PTGES, PTGIS, or TBXAS1 generated from the Maurer et al., dataset representing 124 PDAC patients.

## Methods

### Mice

Mice were housed in accordance with NIH guidelines in AAALAC-accredited facilities at the Salk Institute for Biological Studies. The Salk Institute IACUC approved all animal studies. *LSL-Kras*^*G12D/+*^, *Ptf1a*^*Cre/+*^, *Pdx1Cre*, *Trp53*^*R17H*^, and *Trp53*^*fl/fl*^ mice have previously been described (Bardeesy et al., 2006; Hingorani et al., 2005). C57B6/J mice were either purchased from the Jackson Laboratory (Bar Harbor, ME) or bred in-house. CD1 mice were either purchased from Charles River Laboratories (Wilmington, MA) or bred in-house.

### Orthotopic Transplantation

Cells lines (FC-1199, FC-1242, and FC-1245)(Engle et al., 2019) were cultured in 10% FBS (Biowest) and 1% Anti-Anti (Gibco) in DMEM (Gibco). Medium was replaced every three days and plates were split at a 1:4 ratio when reaching 80% confluency. To prepare for transplantation, cells were dissociated with 0.05% Trypsin (Gibco) for 2-4 minutes, and the enzyme was quenched with culture media. Cells were diluted to 100 cells per 20 μl of 1:1 Matrigel (Corning):medium and placed on ice until transplantation. Mice were administered anaesthesia and analgesics in accordance with IACUC protocol. A patch of skin over the spleen was shaved, disinfected with Nolvasan (Patterson Veterinary Supply), and cut to expose the spleen. The pancreas was exposed via gripping the spleen with circular forceps. One hundred cells resuspended in Matrigel:medium were injected into the pancreas using an insulin syringe. Organs were replaced, the peritoneum was sutured, and the skin was stapled. The mice were allowed to recover and were monitored in accordance with IACUC protocol.

### Human samples

Distribution and use of all human samples was approved by the Institutional Review Boards of the Salk Institute for Biological Studies and Vanderbilt University. Flash frozen human pancreas samples were either purchased from Indivumed (Frederick, MD) or were acquired from the Cooperative Human Tissue Network (CHTN). All normal pancreas samples were acquired from PDAC patients and were pathologically determined to be normal.

### Comprehensive eicosanoid panel

Eicosanoid profiling was conducted on flash-frozen pancreas tissue from wild type mice (C57B6/J or CD1), *KPC* mice, or orthotopic tumors (cell lines FC-1199, FC-1242, and FC-1245) grown in C57B6/J mice. In separate experiments, eicosanoid profiling was performed on flash frozen human PDAC or normal pancreas. Age-matched normal and PanIN-bearing pancreata were included from a previous study (DelGiorno et al., 2020a). Tissues were homogenized in 1 ml of PBS containing 10% ethanol and 300 μl were extracted using strata-x polymeric reverse phase columns (88-S100-UBJ Phenomenex). Samples were taken up in 50 μl of 63% H_2_0, 37% ACN, 0.02% Acetic Acid, and 10 μl was injected into UPLC (ACQUITY UPLC System, Waters) and analyzed on a Sciex 6500 Qtrap mass spectrometer at the UCSD Lipidomics Core as previously described (Quehenberger et al., 2010). Tissue eicosanoid concentrations were quantified using deuterated internal standards in conjunction with standard curves obtained in parallel using identical conditions as previously described (Wang et al., 2014), and were normalized to total protein mass of the sample.

### Histological staining

Tissues were fixed overnight in zinc-containing, neutral-buffered formalin (Fisher Scientific), embedded in paraffin, cut in 5 μm sections, mounted, and stained. Sections were deparaffinized in xylene, rehydrated in a series of graded ethanols, and then washed in PBST and PBS. Endogenous peroxidase activity was blocked with a 1:50 solution of 30% H_2_O_2_:PBS followed by microwave antigen retrieval in 100 mM sodium citrate, pH 6.0. Sections were blocked with 1% bovine serum albumin and 5% normal goat serum in 10 mM Tris (pH 7.4), 100 mM MgCl_2_, and 0.5% Tween-20 for 1hr at room temperature. Primary antibodies (Table S1) were diluted in blocking solution and were incubated on tissue sections overnight. Slides were then washed, incubated in streptavidin-conjugated secondaries (Abcam) and developed with DAB substrate (Vector). Hematoxylin and eosin (H&E) staining was performed to assess tissue morphology. Immunofluorescence on paraffin-embedded tissues followed the immunohistochemistry protocol until the blocking step. Instead, tissues were blocked with 5% normal donkey serum and 1% BSA in 10 mM PBS for 1 hour at room temperature. Tissue sections were stained with primary antibodies in 10 mM PBS supplemented with 1% BSA and 0.1% Triton X-100 overnight. Sections were then washed 3 × 15 min in PBS with 1% Triton X-100, incubated in Alexa Fluor secondary antibodies, washed again for 3 × 5 min, rinsed with distilled water, and mounted with Prolong Gold containing DAPI (Invitrogen). All slides were scanned and imaged on an Olympus VS-200 Virtual Slide Scanning microscope.

### Immunohistological analysis and scoring of patient samples

H&E and immunohistochemistry of PTGES, PTGIS, and TBXAS was conducted on serial tissue sections obtained from 22 treatment naïve PDAC patients (34 slides total) and was then analyzed with QuPath version 0.3.0, an open-source software for digital pathology and whole-slide image (WSI) analysis, as previously described (Apaolaza et al., 2021; Bankhead et al., 2017). Representative H&E WSI were registered with serial sections of PTGIS, PTGES, and TBXAS1 immunostaining using the “Interactive Image Alignment” extension. Up to three 500 μm × 500 μm regions of interest (normal ducts, acinar-to-ductal metaplasia, low-grade PanIN, high-grade PanIN, well-differentiated PDAC, poorly-differentiated PDAC, myxoid fibrosis and compact fibrosis) were selected on the H&E slide, and the same registered area of the immunostaining was scored by two observers, including one board-certified pathologist subspecialized in gastrointestinal diseases including pancreas (V.Q.T). The epithelial and stromal compartments were scored separately for each component. The scoring scheme for the epithelium was: 0=negative or very rare cells, 1=mild staining in less than 50% of cells, 2=intermediate, 3=more than 50% of cells staining strongly. The scoring scheme for the stromal compartment was: 0=negative or very rare cells, 1=few interspaced cells with mild staining, 2=intermediate, 3=frequent approximated cells with strong staining together.

### Tumor dissociation and fluorescence activated cell sorting (FACS)

Orthotopic tumors were harvested near endpoint (4 weeks for FC-1245, 6 weeks for FC-1199), washed twice with DMEM (Gibco), chopped, and dissociated with 200 mg DNASE I (Sigma-Aldrich), 0.2 mg Pronase (Sigma Aldrich), and 1 mg Collagenase P (Sigma Aldrich) in 10 ml Gey’s solution (Sigma Aldrich) for 50 minutes at 37°C. Cells were passed through a 100 μm filter and washed with 2% FBS (Biowest) in HBSS (Gibco). Cells were incubated with 1 ml ACK lysis buffer (Gibco) for 3 minutes on ice and were then washed twice. Cells were resuspended in 500 μm 2% FBS in HBSS and incubated on ice in the dark with 1 μl Fc block (BD Biosciences) for 3 minutes. Cells were then stained with CD45 (APC, Biolegend) and EpCAM (PE/Cy7, Biolegend), resuspended in 2% FBS in HBSS containing 5μg/ml DAPI (Invitrogen), and sorted into DMEM containing 10% FBS. FACS was conducted at the Salk Institute’s Flow Cytometry Core facility on an Aria Fusion cell sorter (100-μm size nozzle, 1 × PBS sheath buffer with sheath pressure set to 20 PSI). Cells were sorted in 1-drop Single Cell sort mode for counting accuracy. EpCAM+;CD45-cells were sorted as the epithelial fraction and EpCAM-CD45+ cells were sorted as the immune cell fraction

### RNA isolation and qRT-PCR

Cells were lysed with 1% beta-mercaptoethanol (Sigma) in RLT plus buffer (Qiagen), and frozen at −80 °C. Samples were thawed, and RNA was isolated with the Qiagen RNeasy Micro Kit (Qiagen, 74004) according to the manufacturer’s instructions. cDNA synthesis was carried out using iScript reagent (Bio-Rad), and RT-qPCR was performed using Power SYBR Green PCR Master Mix (Applied biosystems) on the ABI 7900 detection system (Applied Biosystems). Relative expression values were determined using the standard curve method and were normalized to housekeeping gene Rplp0. Primer sequences can be found in Table S2.

### Bulk Human RNA Sequencing Analysis

Compartment-specific gene expression profiles of human PanIN (n = 26) and PDAC (n = 197) were generated using laser capture microdissection with subsequent RNA sequencing as previously described (Maurer et al., 2019; Maurer and Olive, 2019). Statistics were computed with DESeq2. To study the association of epithelial, stromal or joint eicosanoid enzyme expression, respectively, and patient outcome, we binned our patient cohort (n=197 for epithelial and n=124 for stromal and joint evaluation) into quartiles according to the expression of the respective enzyme and compared differences in outcome between the upper and lower quartile using a log-rank test as implemented in logrank_test function from the lifelines Python package (Davidson-Pilon et al., 2021).

### Single Cell RNA Sequencing Analysis

Processed count matrices for scRNA-seq datasets were downloaded from the Gene Expression Omnibus (GEO) database (Elyada et al., 2019; Peng et al., 2019; Schlesinger et al., 2020). Quality controls for read counts, genes expressed, and mitochondrial gene expression are described in the respective publications. Normalization, variable feature selection, and scaling were performed with Seurat (Stuart et al., 2019). Data preprocessing and dimensionality reduction was performed with Seurat and UMAP coordinates were generated using first 50 components returned by PCA. Cell types were determined in the Peng dataset using annotations provided by the authors and in the Schlesinger and Elyada datasets using panels of gene markers described in the results (Figure S1).

### Analysis of TCGA database

Clinical data associated with the TCGA PAAD database was queried using the cBioPortal website (Cerami et al., 2012; Gao et al., 2013). First, misclassified and non-PDAC patient samples were excluded. Survival analysis was performed between groups representing the top and bottom 25% of samples by expression of a given gene. Statistics were computed using the log rank test.

### Statistical analysis

Statistical analyses, data processing, heatmap plotting, hierarchical clustering, and PCA analysis were performed in R (https://www.r-project.org/) and/or Prism (GraphPad). Statistical significance was calculated by either two-tailed unpaired t-tests assuming equal variance or one-way ANOVA. qRT-PCR and IHC quantification data are expressed as mean ± standard deviation. Eicosanoid levels are expressed as mean ± standard error of the mean.

---

**Table S1.**
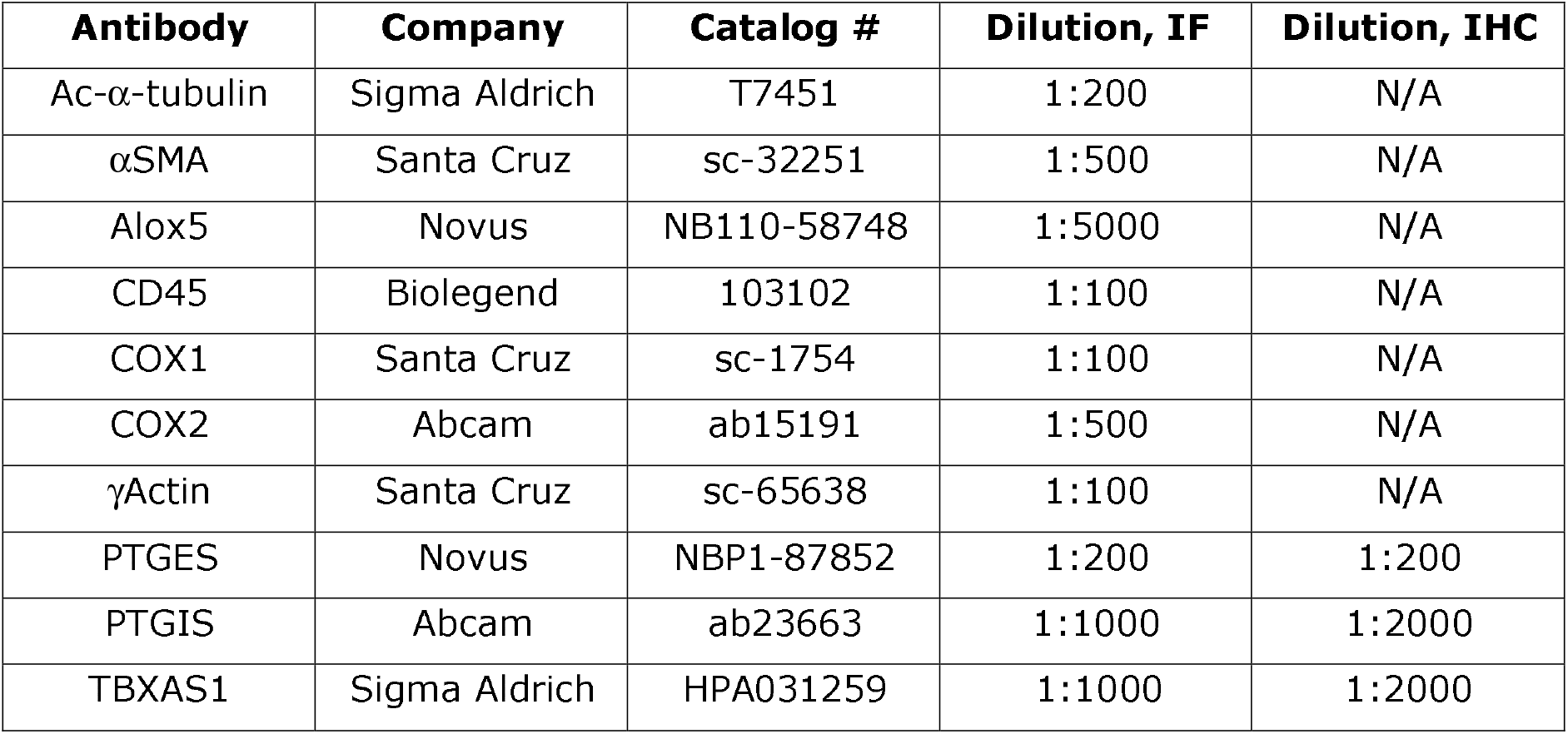
Primary antibodies used in immunofluorescence (IF) and immunohistochemistry (IHC) studies.

**Table S2.**
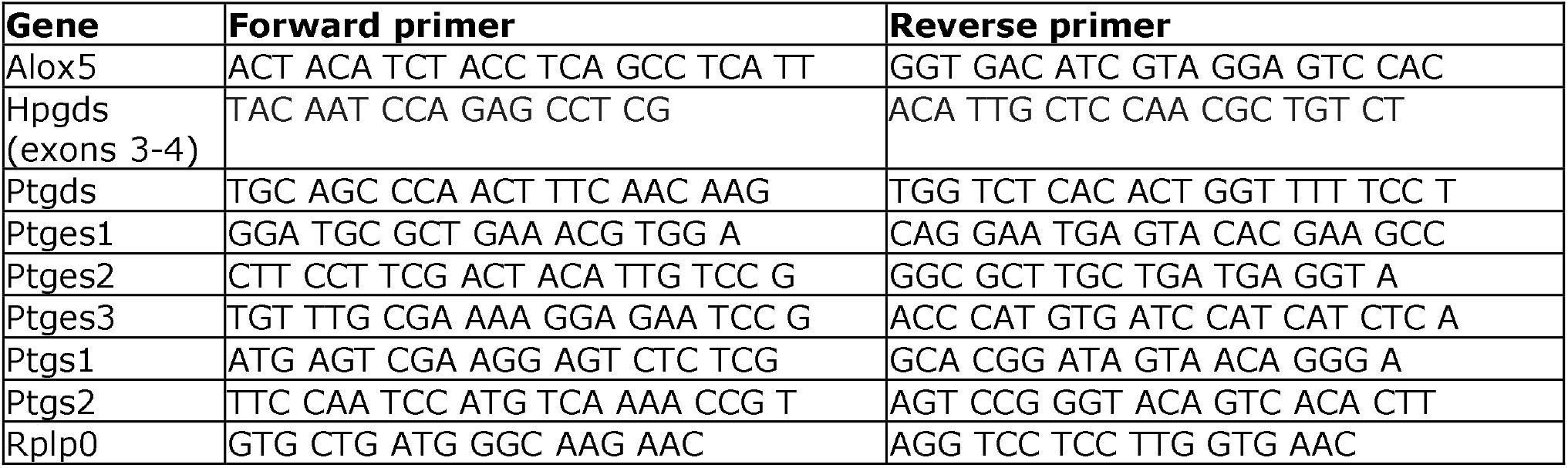
Primers used in qRT-PCR studies.

